# Tri-Functional CRISPR Screen Reveals Overexpression of *QDR2* and *QDR3* Transporters Increase Fumaric Acid Production in *Kluyveromyces marxianus*

**DOI:** 10.1101/2025.09.03.674080

**Authors:** Mackenzie Thornbury, Raha Parvizi Omran, Lalit Kumar, Adrien Knoops, Raghad Abushahin, Malcolm Whiteway, Vincent J.J. Martin

## Abstract

Organic acids such as fumaric acid are widely used in the food and beverage industry as acidulants and preservatives, while also serving as versatile precursors for industrially relevant compounds. Fumaric acid is still predominantly produced through petroleum-derived processes. To enhance production efficiency and diversify supply, we are engineering *Kluyveromyces marxianus* as a biosynthetic platform from renewable feedstocks. In previous work, we have established *K. marxianus* Y-1190 as a host for lactose valorization based on its high growth rate on lactose and its tolerance for acid conditions. Here, we establish a trifunctional genome-wide library for *K. marxianus* using CRISPR activation, interference, and deletion to allow identification of gene expression perturbations that enhance tolerance to fumaric acid. We determined that deletion of *ATP7*, encoding a subunit of the mitochondrial F_1_F_0_ ATP synthase, and overexpression of *QDR2* and *QDR3*, two previously uncharacterized members of the 12-spanner H⁺ antiporter (DHA1) family in *K. marxianus,* can enhance fumaric acid tolerance. We also found that integrated overexpression of both *QDR2* and *QDR3* in a Δ*FUM1* background strain improved titers of fumaric acid production from 0.26 g L^−1^ to 2.16 g L^−1^. Together, these results highlight roles for membrane transport and mitochondrial function in enabling fumaric acid tolerance and production in *K. marxianus*.

**HIGHLIGHTS:**

- Trifunctional CRISPR-AID enables gene activation, interference & deletion in *K. marxianus*.
- Genome-wide CRISPR-AID screen identifies guides conferring fumaric acid tolerance.
- Overexpressing of *QDR2* and *QDR3* increases fumaric acid tolerance and production.
- Deletion of mitochondrial gene *ATP7* significantly improves fumaric acid tolerance.
- Fermentation of *QDR2* and *QDR3* overexpression strain yields 2.16 g L⁻¹ fumaric acid.

## 1. Introduction

Organic acids have a long history of production through fermentation. Citric acid is primarily produced by the filamentous fungus *Aspergillus niger*, while lactic acid is typically produced by *Lactobacillus* species such as *Lb. amylovorus*, the fungus *Rhizopus oryzae*, and recombinant yeast strains (Abedi et al., 2020; FoodIngredientsFirst, 2010; Sauer et al., 2008). However, many other organic acids are not currently manufactured by fermentation at large scale due to high costs and/or low yields. Malic acid, fumaric acid, and adipic acid are examples of building blocks for plasticizers and polymers such as nylon 66 (Ilica et al., 2019; Roa Engel et al., 2008; Van De Vyver and Román-Leshkov, 2013; Q. Xu et al., 2012), and research is ongoing to develop lower-cost fermentation methods for these acids. These approaches include using alternative feedstocks (Liu et al., 2013; Thakker et al., 2013; Zheng et al., 2012) and non-conventional production organisms with inherent qualities that result in higher yields (Jiang et al., 2013; Ju et al., 2020; Pyne et al., 2023; Zhang et al., 2012).

*Kluyveromyces marxianus* is a Crabtree-negative member of the *Saccharomycetaceae* family with many characteristics suited to biomanufacturing. It can metabolize diverse carbon sources (Nonklang et al., 2008), grows rapidly (Groeneveld et al., 2009), and is both heat- and acid-tolerant (Anderson et al., 1986; Chang et al., 2014). Recent advances in genetic engineering methods for *K. marxianus* such as the development of synthetic biology toolkits (Nambu-Nishida et al., 2017; Rajkumar et al., 2019; Thornbury et al., 2025) and CRISPR genome editing (Cernak et al., 2018; Juergens et al., 2018; Lee et al., 2018; Wang et al., 2024; Zhou et al., 2024) have enhanced interest in using this host in biomanufacturing (Goshima et al., 2013; Li et al., 2021; Löbs et al., 2018). The majority of this bioprocess development is done under glucose-fed growth conditions, with only a few studies leveraging *K. marxianus’*s substrate-utilizing diversity by using inulin or sucrose as a carbon source (Rouwenhorst et al., 1988; Zhang et al., 2019).

Lactose, another potential carbon source, is a prominent by-product of the dairy processing industry, and considerable research has focused on valorizing lactose-rich whey and milk permeate (Christensen et al., 2011; Ryan and Walsh, 2016). Within the *K. marxianus* species, dairy strains utilize lactose more efficiently than non-dairy strains due to 13 amino acid variations in the lactose transporter Lac12p (Varela et al., 2017). We have previously characterized *K. marxianus* Y-1190 and found the variations in Lac12p associated with lactose utilization, identifying it as a promising candidate for lactose valorization (Thornbury et al., 2025). This study investigates using lactose from dairy permeate as a feedstock for fumaric acid production by *K. marxianus*.

Fumaric acid is an attractive target for biomanufacturing due to its wide range of industrial applications and its potential for sustainable production. It is widely used in the food and beverage industry, both as an acidulant and in the production of unsaturated polyester resins (Martin-Dominguez et al., 2018). Fumaric acid is currently derived from petroleum-based processes therefore, bio-production offers a renewable alternative that reduces reliance on fossil resources (Werpy and Petersen, 2004). Developing efficient microbial platforms for fumaric acid production will enable cost-effective and scalable bioprocesses, further driving its adoption as a sustainable chemical.

Previous fungal and yeast production of fumaric acid was achieved in *Saccharomyces cerevisiae*, *Rhizopus oryzae*, *Scheffersomyces stipitis*, and *Candida glabrata* (Chen et al., 2019, 2016; Huang et al., 2010; Wei et al., 2015)*. R. oryzae* is the most prolific producer reaching a final titer of up to 103 g L^-1^ in 20 L fermentation tanks (Rhodes et al., 1962). As alternatives to *R. oryzae*, recombinant strains were developed by expression of the reductive TCA genes of *R. oryzae* in *C. glabrata* and *S. cerevisiae* to achieve titers of 21.6 g L^−1^ and 33.13 g L^−1^, respectively (Chen et al., 2019, 2016). *Escherichia coli* studies have focused on intensive rewiring of central metabolism to achieve titers of 42.5 g L^−1^ (N. Li et al., 2014). While these efforts have demonstrated the potential for high-titer production, challenges related to acid tolerance and utilization of substrates other than glucose highlight the need for alternative hosts such as *K. marxianus*.

In the present study, we deleted the fumarate hydratase *FUM1* gene in *K. marxianus* to increase fumaric acid titers. While *FUM1* deletion promoted fumaric acid accumulation, it also impaired acid tolerance. To address this trade-off, we used CRISPR-AID selection to uncover genetic perturbations that could restore growth under acid stress. This study adapts CRISPR-AID for use in *K. marxianus* and identifies candidate genes that support both acid tolerance and fumaric acid production. The findings suggest that the proteins Qdr2p and Qdr3p likely function as fumaric acid exporters, mitigating intracellular stress and increasing fumaric acid production. This work provides valuable insights into engineering *K. marxianus* for organic acid production and industrial biotechnology applications.

## 2. Methods

### 2.1. Plasmids, Strains and Growth Media

*Escherichia coli* strain DH5α, used to maintain and amplify plasmids, was cultured at 37 °C in Luria broth containing the appropriate antibiotics. *K. marxianus* strain Y-1190 and Y-1190Δ*FUM1* were used as the host for all genome engineering in this work. Unless otherwise stated, *K. marxianus* was cultivated and maintained in YPD (10 g L^−1^ Bacto Yeast Extract, 20 g L^−1^ Bacto peptone, 20 g L^−1^ glucose). Plasmids, oligonucleotides, and engineered strains used in this study can be found in Table S1, Table S2, and Table S3, respectively.

### 2.2. *K. marxianus* Transformations and CRISPR-Cas9 Editing

All sequences of *K. marxianus* Y-1190 were obtained from genome assembly ASM4656278v1. Unless otherwise stated, *K. marxianus* genetic modifications were facilitated using double-sgRNA CRISPR-Cas9-mediated deletions and integrations (Thornbury et al., 2025). The SpCas9 and sgRNAs were delivered by pMTKM01 (Addgene plasmid #233473) and edited strains were selected on YPD agar containing 100 µg mL^-1^ nourseothricin.

Transformations of *K. marxianus* were performed according to a previously published protocol with the following modifications (Antunes et al., 2000). In brief, 1 mL of *K. marxianus* cells in exponential phase were centrifuged at 1860 g for 3 min. After washing once with 1 mL of sterile H_2_O, the cells were suspended in 10 µL of 10 mg mL^-1^ salmon sperm, and 500 ng of sgRNA plasmid was added, along with 1 µg of linearized donor DNA for genome integration. To the cells and DNA, 400 µl of transformation buffer (40 % polyethylene glycol 3350, 0.1 M lithium acetate, 10 mM Tris–HCl (pH 7.5), 1 mM EDTA, and 10 mM DTT) was added to a 1.5 mL microfuge tube and the solution was mixed by agitation. The transformation mix was then incubated at room temperature for 15 min followed by heat shock at 47 °C for 15 min. The cells were then centrifuged at 1860 g for 3 min and suspended in 400 µl of YPD to recover overnight at 30 °C without shaking. The next day the cells were plated on YPD agar with appropriate selection. If necessary, an outgrowth step was done where 50 µl of recovered transformation was grown in 500 µl of YPD with selection for 2-3 days. This outgrowth was diluted 1:1000-10000 before plating on YPD agar with selection.

### 2.3. Establishing CRISPR-AID in *K. marxianus*

CRISPR-AID (Activation, Interference, Deletion) is a tri-functional platform that enables simultaneous gene activation, repression, and deletion, providing a powerful tool for functional genomics and metabolic engineering, first built in *S. cerevisiae* to enhance furfural tolerance (Lian et al., 2017). To adapt and test the efficiency of CRISPR-AID in *K. marxianus,* the mNeonGreen (mNG) CDS was inserted at the *URA3* locus using CRISPR-Cas9 allowing *URA3* auxotrophy for sgRNA plasmid selections while providing a measurable phenotype. This strain (MTK019) also expressed the three CRISPR-AID Cas enzymes (Lian et al., 2019); however, we replaced the promoters driving Cas expression with *K. marxianus* promoters, and replaced the terminators with either *K. marxianus* or *S. cerevisiae* terminators known to be active in *K. marxianus*. The resulting plasmid (pMT195) was integrated into the I3 locus between *KmSWF1* and *KmARO1* (Rajkumar et al., 2019) using CRISPR-Cas9. Guides for activation, interference and deletion of the mNG cassette were designed on CCTop (Stemmer et al., 2015) and expressed from plasmids pMT149-pMT156. These plasmids were transformed into MTK019 and three clones per sgRNA were picked into SC-URA media containing G418. Cultures were grown overnight and then back diluted to OD_600_ of 0.05 and grown for an additional 24 h. The cultures were diluted 1:10 and fluorescence was measured at excitation 510 nm and emission 545 nm in a plate reader (Infinite200, Tecan).

### 2.4. Genome-Wide CRISPR-AID sgRNA Library Design and Construction

#### 2.4.1. Building the CRISPR-AID Strain

To integrate the CRISPR-AID cassette into *K. marxianus* Y-1190Δ*FUM1*, we used a similar method as described above, however, we replaced the constitutive promoters with tetracycline inducible promoters resulting in plasmid pAK104. Other than the *Cas* coding sequences, all parts were sourced from the following tool kits: Lee *et al*., 2015, Rajkumar *et al*., 2019, and Thornbury *et al*., 2025. The three *Cas* sequences and their scaffolds were amplified from CRISPR-AID plasmids from Lian *et al*. 2019 (Addgene (Lian et al., 2019) #136989, #136992-136994). A total of 5 ug of pAK104 was digested with NotI-HF (New England Biolabs) and the linear plasmid used as a donor. We used CCTop to generate sgRNAs targeting the I3 site between *SWF1* and *ARO1*, and selected sgRNAs displaying a CRISPRater score of 0.60 or higher (Labuhn et al., 2017). We used Golden-Gate cloning as previously described to introduce two sgRNAs into the pMTKM01, a SpCas9 vector (Dykstra et al., 2023; Engler et al., 2008; Thornbury et al., 2025).

#### 2.4.2. CRISPR-AID sgRNA Library Design

Genome-wide library sgRNA sequences were designed using CHOPCHOP via the command line (Labun et al., 2019, 2016; Montague et al., 2014). Promoter sequences were designated as 400 bp upstream of each transcriptional start site and were used to design activation and interference sgRNAs, while coding sequences were used for the deletion sgRNAs. *K. marxianus* preferentially uses non-homologous end joining (NHEJ) for DNA repair (Abdel-Banat et al., 2010), and therefore we did not design donor DNA for deletions. As proof of concept, we demonstrated that a single sgRNA was sufficient to disrupt the mNG reporter signal (Fig 1A), validating our approach.

**Figure 1.**
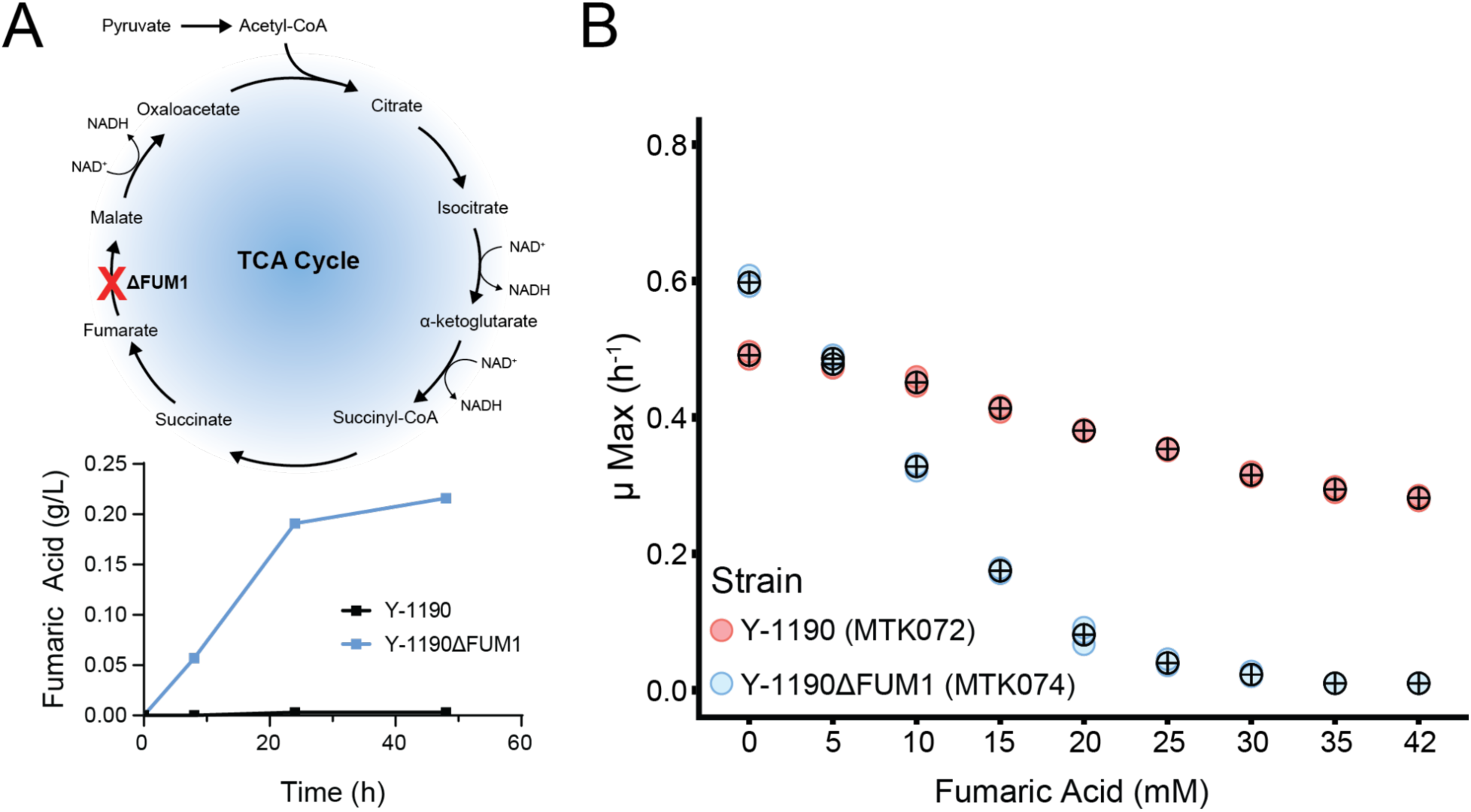
Effect of *FUM1* deletion on fumaric acid production and tolerance **A**. Schematic of the TCA cycle and the *FUM1* deletion that enhanced fumaric acid production in *K. marxianu*s. **B**. Growth rate of *K. marxianu*s CRISPR-AID strain with (MTK074) and without (MTK072) *FUM1* deletion. Black circles are the mean of three biological replicates.

Because CHOPCHOP generates all possible sgRNA sequences for a given DNA target sequence, we adapted a previously published ranking criterion to select sgRNAs likely to be the most efficient (Lian et al., 2019) (Table S4). We selected the top six sgRNAs for activation and interference, and the top four for deletion. In rare cases, no PAM sequence was available within the selected target region, or no unique sgRNAs could be designed resulting in less than four, or six, sgRNAs being ordered for that gene. Summary coverage of our library can be found in Supplementary Table S5. The final three libraries consisted of 26185 activation, 26988 interference and 18763 deletion sgRNAs; all sequences can be found in Table S5. When combined, the library contains a total of 71,936 guides covering 4,740 of the 4,853 annotated genes. The remaining 113 genes correspond to transposable elements with significant sequence overlap, which prevented unique guide design. While we aimed to include six guides per gene for activation and interference, limitations such as a lack of unique PAM sites or overlapping promoter regions prevented full coverage: For activation and interference, 83% and 86% of genes, respectively, had full six guide coverage, and 98.5% and 99.6% of genes were covered by at least one guide (Table 1). For deletion, 98% of genes had complete four-guide coverage, and 99.5 % were targeted by at least one guide. Adapters were added to both ends to facilitate PCR amplification that corresponded to the flanking tRNA_glycine_ sequence within the plasmid, allowing all three sgRNA types to be amplified with one set of primers. All sgRNA libraries were synthesized by Twist Bioscience.

**Table 1.** Summary Coverage of the *K. marxianus* CRISPR-AID Library.

|  | Number of Genes Targeted |  |  |  |  |  | Total Genes | Total sgRNAs |
| --- | --- | --- | --- | --- | --- | --- | --- | --- |
| # of sgRNAs | 1 | 2 | 3 | 4 | 5 | 6 |  |  |
| #of genes in ActLib | 97 | 118 | 132 | 165 | 218 | 3951 | 4740 | 26185 |
| #of genes in IntLib | 29 | 74 | 105 | 167 | 236 | 4108 | 4740 | 26988 |
| #of genes in DelLib | 14 | 25 | 21 | 4659 | 0 | 0 | 4740 | 18763 |

#### 2.4.3. sgRNA Library Construction

The pooled oligos from each of the three libraries were suspended in TE buffer to a final concentration of 10 ng μL^-1^. An initial 50 μL Phusion high fidelity short PCR amplification of 1 ng of each library was done using blunt primers (MT736 & MT737). Because of the different sizes of oligos to be amplified, different amounts of amplification cycles were done. Activation sgRNAs for LbCas12a have shorter scaffolds and were amplified for 9 cycles, whereas SaCas9 and SpCas9 scaffolds are similar in length, as such, deletion and interference sgRNAs were amplified for 12 cycles. PCR amplicons were purified by kit with an isopropanol addition for small fragments (GeneJet, Thermofisher). The purified amplicons were used as template for a second PCR to add Golden Gate adapters for downstream cloning. Twelve 50 μl Phusion high fidelity polymerase PCR reactions were done per library with 10 ng of template and with PCR cycles kept to 9-12 as before. The 12 reactions were pooled, and gel purified (GeneJet, Thermofisher) using a 0.8 % agarose gel.

The purified sgRNA libraries were cloned into a GFP-dropout destination vector containing a *SNR52* promoter and a *SUP4* terminator (pMT351) using the Golden Gate cloning method (Engler et al., 2008). Briefly, 20 fmol of pMT351 and 40 fmol of each library were added to PCR tube with 10 U BsaI-HF, 200 U T4 ligase, 1.5 μl T4 ligase buffer, 1.5 μl BSA/PEG mix (1 mg mL^-1^ BSA, 10 % PEG 3350) and ddH_2_O added to a final volume of 15 μl. The libraries were ligated using a thermocycler and 45 cycles of 5 min 37 °C digest and 16 °C ligate with a final 1-h 37 °C digestion step. The Golden Gate ligations were purified by isopropanol DNA purification (Green and Sambrook, 2017) and 200 ng of each library was transformed into 40 μl of *E.coli* MegaX DH10B T1^R^ Electrocomp Cells (Thermofisher). Immediately after electroporation, 1 mL of pre-warmed SOC media was added before transferring the cells to a 15 mL conical tube containing 3 mL of pre-warmed SOC. The transformed library was recovered at 37 °C shaking for 1 h before plating on two 24.5 cm x 24.5 cm bioassay plates of LB-Carbenicillin. An aliquot of 50 μl was retained for serial dilutions to count colonies. We achieved over 10-fold transformation coverage for each library (Table S6). For each library, 25 mL of cold LB was used to scrape each bioassay plate resulting in ∼45 mL of transformed *E. coli* culture per library, which was divided between two maxi prep columns (GeneJet, Thermofisher) for DNA extraction.

Construction of our CRISPR-AID sgRNA libraries in *K. marxianus* was done as previously described in Lian et al., (2019) for *S. cerevisiae* with the following modifications. For each library, five transformations with 10 μg of DNA were done according to the protocol defined above. All five transformations were combined in 4 mL of YPD and recovered at 30 °C overnight. The next morning, the recovered cultures were topped up to 10 mL in YPD and 5 mL each were plated on SC-URA bioassay plates. Fifty μl were set aside to dilute 1:100 and 1:1000 for colony counting. After determining colony count for each library, the libraries were scraped into cold SC-URA and pooled to ensure equal representation. Individual and pooled libraries were adjusted to 20 % total glycerol and stored at -80 °C.

### 2.5. CRISPR-AID Selection in Fumaric Acid

A fumaric acid growth inhibition pilot experiment was done that identified 25 mM of fumaric acid as our starting concentration to select CRISPR-AID library clones (Fig. S2). For identifying clones with increased tolerance to fumaric acid, the pooled library was inoculated at an OD_600_ of 0.1 in 50 mL of SC-URA with anhydrous tetracycline (aTC) at 500 ng μl^-1^ with either 0 mM or 25 mM fumaric acid and grown in a shaking incubator at 30 °C and 220 rpm. Cultures were sampled when growth in the 25 mM condition had exceeded that of the control (EV, plasmid with no sgRNA). To increase the screening stringency, cultures selected on 25 mM fumaric acid were back-diluted to an OD_600_ of 0.1 into the same medium supplemented with 30 mM fumaric acid, grown for an additional 24 hr and harvested. This back dilution step was repeated once more with a medium supplemented with 42 mM fumaric acid and a 30-hr growth period. The control cultures did not grow in 30 mM and 42 mM fumaric acid conditions.

### 2.6. CRISPR-AID sgRNA Enrichment Analysis

Cells from each library were harvested at an OD_600_ unit of 0.5, corresponding to ∼3.0×10^7^cells, by centrifugation (1860 g for 3 min) and sgRNA plasmids were extracted from the cells using a Zymoprep yeast miniprep kit (Zymo Research). sgRNAs were amplified using Platinum SuperFi DNA polymerase (Invitrogen, 12351010) and primers containing heterogeneity spacers for 20 cycles from ∼8 ng of plasmid template (Table S2). Following amplification, two 50 μl, 20 cycles, PCR reactions were performed per sample using Phusion high fidelity (New England Biolabs) and primers with Nextera index adapter (Illumina) sequences. The PCR products were purified (GeneJet, Thermofisher) and 130 ng of each sample were pooled. The pooled amplicons were sequenced on an Illumina NextSeq platform using 300 paired-end reads (Génome Québec).

Initial quality control was done using FastQC v0.11.5 and MultiQC v1.12 (Cock et al., 2010; Ewels et al., 2016). The paired reads were merged using PEAR v0.9.6 with a minimum read length (-t) of 20-bp and a quality score threshold (-q) of 25 (Zhang et al., 2014). We were able to merge 99.5 % of reads, for an average of 1.1×10^6^ merged reads per condition (Table S7). We were able to maintain a minimum of 10-fold theoretical coverage for each condition, except for replicate B for the 42 mM fumaric acid condition. This replicate was included in downstream analysis despite lower coverage. These merged reads were binned into activation, interference, or deletion sgRNAs using the scaffold sequences and Cutadapt v5.0 (Martin, 2011). Binned sgRNA sequence counts were analyzed using MAGeCK v0.5.8 (W. Li et al., 2014). sgRNA enrichment was measured over the 0 mM fumaric acid control condition. Targets with the highest enriched sgRNAs that had significantly low p-values and false discovery rates were selected for further testing.

### 2.7. Validation of CRISPR AID Hits

The top 10 over-represented sgRNAs from the library screen were all activation sgRNAs. To validate them as true hits, we overexpressed in *K. marxianus* Y-1190Δ*FUM1* the target gene that corresponded to the enriched sgRNA and measured strain fitness in a medium supplemented with fumaric acid. The genes were amplified from *K. marxianus* Y-1190 genomic DNA and cloned into a vector with the strong promoter *KmNC1p* (Addgene plasmid #233500). As *K. marxianus* Y-1190 is a heterologous diploid, the PCR reaction had a chance of amplifying both the primary and the haplotig allele. In these cases, we included both in the validation screen.

### 2.8. Growth Rate Analysis

Three independent colonies of each transformed strain were inoculated into 96-well deep plate containing 700 μl of Synthetic Complete Lactose (SC-Lactose; 6.7 g L^−1^ yeast nitrogen base with ammonium sulfate, 20 g L^−1^ lactose, 1.92 g L^−1^ yeast synthetic drop-out medium supplement without uracil, 0.08 g L^−1^ uracil) media supplemented with 200 μg mL^-1^ G418. To test strain fitness, cell growth was measured from overnight cultures back diluted to an OD_600_ of 0.1 in the same medium but with concentrations of fumaric acid varying from 0 mM to the solubility limit of 42 mM. Cultures were incubated at 30 °C in clear, flat-bottomed 96-well microtiter plates and wrapped with Parafilm to minimize evaporation. Absorbance readings were taken at 595 nm with a Sunrise absorbance microplate reader (Tecan) every 20 min over the course of up to 3 days. Growth rates (μ_max_) were calculated in R using the “growthrates” package (Petzoldt, 2022).

### 2.9. Metabolite Analysis

To assess the effect of increased strain fitness on fumaric acid production we measured substrate uptake and metabolites produced by our strains. Shake-flask cultures for fumaric acid production quantification were grown in SC-Lactose or 0.5x supplemented dairy permeate (Thornbury et al., 2025). Following overnight growth in SC-Lactose, cultures were diluted to an OD_600_ of 0.2 in 20 mL SC-Lactose in a 125 mL baffled Erlenmeyer flask. Cultures were incubated at 30 °C, and shaking at 200 rpm for 72 h. At 24 h intervals, 250 μl of the culture supernatant was sampled and frozen at -20 °C. For HPLC analysis, supernatant samples were diluted 1:10 in distilled H_2_O. Metabolites were measured using an Agilent 1290 Infinity II HPLC equipped with an Aminex HPX-87H column (Bio-Rad). Metabolites were resolved isocratically in 10 mM sulfuric acid at 65 °C at a flow rate of 0.6 ml min^-1^. A UV-visible detector was used for the quantification of fumaric acid (260 nM), and a refractive index detector was used to quantify lactose. All measurements were compared to a linear standard curve and done in duplicate or triplicate.

### 2.10. Fed-Batch Cultivation

Controlled fed-batch fermentations were performed in 3 L BioBundle bioreactors (Applikon) at 30 °C. Batch fermentations were initially carried out without pH maintenance for the first 24 h. Thereafter, the pH was maintained at 4.0 by automated addition of 4 N NaOH throughout both batch and fed batch fermentations. Dissolved oxygen was maintained at 20% of air saturation by automatically adjusting the agitation speed (400-600 rpm) and aeration rate (500-600 mL min⁻¹). Off-gas O₂ and CO₂ concentrations were monitored using a Tandem Multiplex Gas Analyzer (Magellan BioTech). Bioreactor inoculums were prepared in two 125 mL Erlenmeyer flasks containing 25 mL batch medium of 0.5x supplemented permeate, grown for 48 h at 30 °C and shaking at 200 rpm. Cultures were used to inoculate at OD₆₀₀ = ∼0.1 in 1 L of batch medium (50 g Lactose, 8.361 g urea, 2.065 g KH₂PO₄, 1.15 g MgSO₄·7H₂O, 1 g NiSO₄·6H₂O, 5 mL vitamin stock, and 5 mL trace element solution per liter). The cultures were grown in batch mode until lactose depletion (49 h), followed by fed-batch operation initiated using a four-fold ultrafiltration-concentrated permeate (Agropur Cooperative) containing 200 g L^-1^ lactose with 33.44 g urea, 8.26 g KH₂PO₄, 4.6 g MgSO₄·7H₂O, 4 g NiSO₄·6H₂O, 20 mL vitamin stock, and 20 mL trace element solution per liter. Samples were collected every 24 h for OD₆₀₀ and HPLC analysis for a total of 7 days. Biomass concentration (g L^-1^) was calculated using a gravimetrically determined conversion factor of 0.22 g L^-1^ per OD₆₀₀ unit. Process data were recorded and analyzed using BioXpert V2 (Applikon). Fermentation samples were diluted 1:5 to 1:20 with deionized water and analyzed by HPLC as described above. Fermentation profiles of dissolved oxygen, feed pump, and pH can be found in Figure S5.

### 2.11. Protein Alignments

Alignment of proteins Qdr2 and Qdr3 from *K. marxianus* Y-1190 were done using Clustal Omega (Madeira et al., 2024) and visualized using Jalview (Waterhouse et al., 2009). Summary alignments were done using NCBI protein blast and transmembrane domains were predicted using DeepTMHMM (Hallgren et al., 2022).

## 3. Results

### 3.1. Creation of a Fumaric Acid Producing *K. marxianus*

To develop a fumaric acid production strain, we disrupted the TCA cycle by deleting *FUM1*, which encodes fumarate hydratase. This strategy was shown to enhance fumaric acid titers in *S. cerevisiae* and *E. coli* (Song et al., 2013; G. Xu et al., 2012b). In *K. marxianus*, *FUM1* deletion similarly increased fumaric titers from 0 to 0.22 g L^-1^ but also compromised the strain’s growth in fumaric acid (Fig. 1). The fumaric acid concentration-dependent growth defect observed in the *FUM1* deletion strain suggests an acid stress phenotype (Fig. 1B). To restore acid tolerance, we designed and implemented a CRISPR-AID system to identify gene expression perturbations that enhance growth under fumaric acid stress.

3.2. Validation of CRISPR-AID in *K. marxianus*

To assess CRISPR-based gene activation, interference, and deletion in *K. marxianus,* we measured GFP fluorescence of *K. marxianus* expressing the three CRISPR-AID Cas proteins, mNG and sgRNAs targeting mNG and its promoter region (MTK019). Two of the three sgRNAs (A1 and A3) designed for activation exhibited 4.6- and 2.6-fold higher fluorescence compared to the wild-type and control strains, indicating successful transcriptional activation (Fig. 2A). All interference sgRNAs (I1, I2 and I3) resulted in a 2.9-to-5.7-fold decrease of fluorescence, and both deletion sgRNAs (D1 and D2) resulted in complete elimination of fluorescence, demonstrating effective repression of gene expression from interference and deletion, respectively (Fig. 2A).

**Figure 2.**
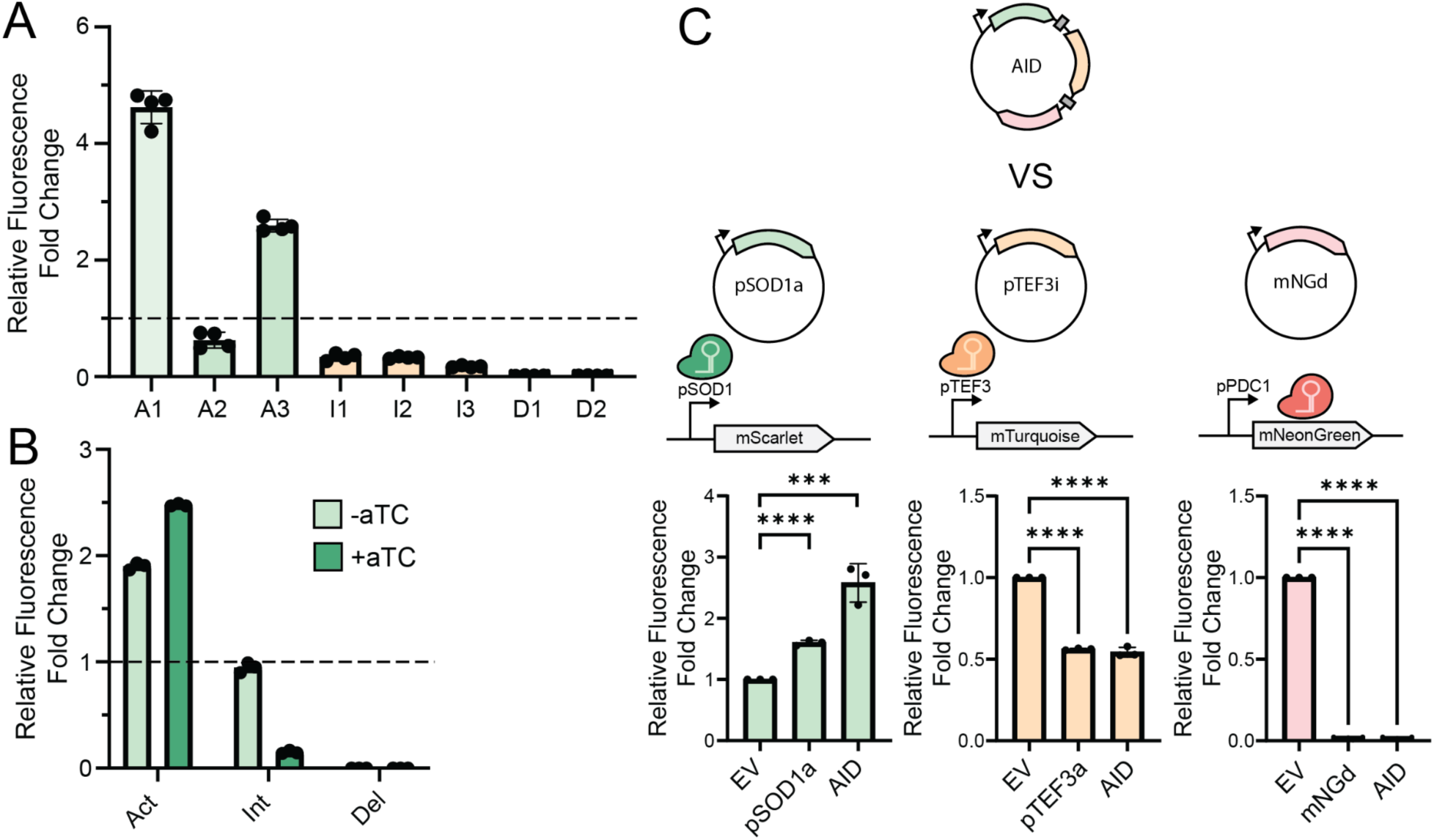
CRISPR-AID functionality in *K. marxianu*s **A**. Constitutive 3 Cas system integrated into Y-1190 transformed with sgRNA plasmids for each perturbation to mNG **B**. CRISPR-AID system with inducible promoters integrated into Y-1190 and transformed with sgRNAs for mNG with or without aTC inducer **C**. Single and combinatorial CRISPR-AID sgRNAs for activation of mScarlet, interference of mTurquoise, and deletion of mNG. Each black dot represents a biological replicate and error bars represent standard deviation.

High plasmid transformation efficiency is critical for constructing a genome-wide CRISPR library in yeast. We observed low transformation efficiency in MTK019, likely due to the constitutive expression of the three Cas proteins causing a burden on the cells (Fig. S1A). To address this, we used a aTC inducible version of the three Cas genes (MTK072) (Thornbury et al., 2025). To ensure that the CRISPR-AID system maintained its function, we transformed the modified stain with the previously validated A1, I1, and D1 sgRNA plasmids and measured GFP fluorescence. We observed an increase in fluorescence with the activation sgRNA, and complete repression of fluorescence with the deletion sgRNA irrespective of addition of the inducer suggesting some leakiness in the inducible promoters (Fig. 2B). For the interference sgRNA, we only observed reduced fluorescence in the presence of aTC suggesting tighter regulation of dSpCas9-RD1152. Despite the leaky promoters for dLbCas12a-VP and SaCas9, the transformation efficiency of MTK072 increased by 5.4-fold ensuring adequate coverage of our sgRNA library (Fig. S1A).

After validating the CRISPR-AID system in *K. marxianus*, we explored the potential for combinatorial perturbations by constructing arrayed sgRNA plasmids targeting multiple genes for simultaneous activation, interference, and deletion. While this combinatorial strategy was not applied in this study, our proof-of-concept demonstrates its feasibility and potential as a powerful tool for multiplexed strain engineering in non-conventional yeasts. To test this system, we modified the MTK019 strain to express two additional reporter fluorescent proteins, mTurquoise and mScarlet, generating MTK063. sgRNAs designed to activate mScarlet, interfere with mTurquoise expression, and delete mNG were expressed in MTK063 individually or in combination from a single plasmid and fluorescence of all three proteins was measured (Fig. 2C). Both single-sgRNA and combination-sgRNA expression resulted in the expected genetic perturbation: mScarlet fluorescence increased 1.6-2.6-fold, mTurquoise decreased 1.8-1.9-fold, and mNG fluorescence was eliminated (Fig. 2C). Notably, single-sgRNA activation constructs produced a moderate 1.6-fold increase in fluorescence over the control (EV) but showed enhanced 2.6-fold activation when expressed in the combinatorial plasmid. In contrast, interference and deletion strategies remained consistent between single- and combinatorial-sgRNA plasmids (Fig. 2C). Together, these data confirm the effectiveness of CRISPR-based activation, interference, and deletion strategies in regulating gene expression in *K. marxianus*.

### 3.3. CRISPR-AID Library Screen

Enrichment of the Δ*FUM1* strain for increased tolerance to fumaric acid was initiated in 25 mM, a concentration determined through a pilot experiment, with 0 mM fumaric acid as the control (Fig. S2). After 144 hr, growth of the CRISPR-AID library surpassed the growth of the control strain. The culture was iteratively back-diluted into 30 mM and subsequently 42 mM fumaric acid, reaching the aqueous solubility limit of fumaric acid. At each concentration, samples were collected for Illumina sequencing to assess sgRNA enrichment (Fig. 3A).

**Figure 3.**
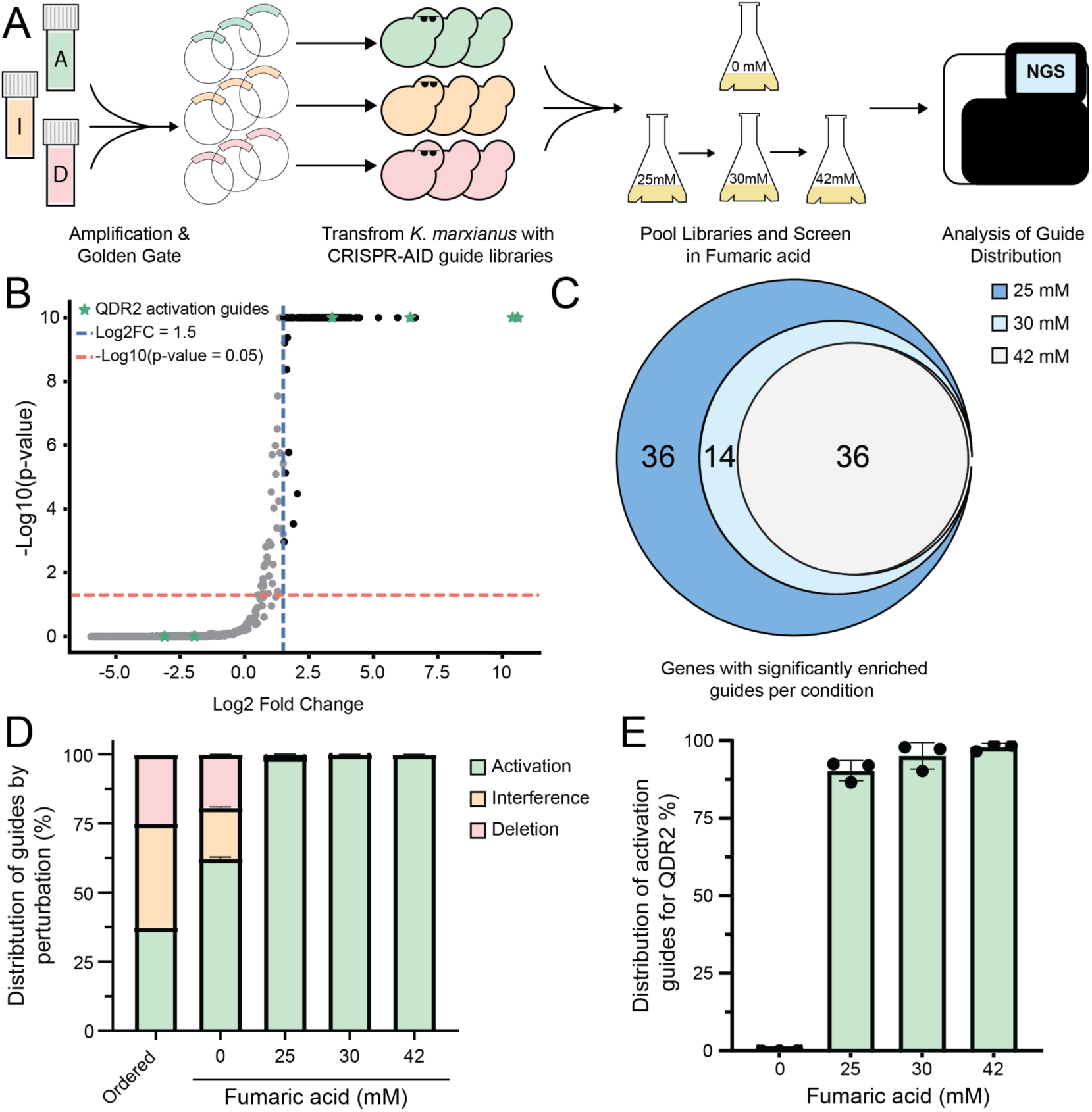
Library workflow and summary statistics **A**. Schematic of library building and screening workflow. **B**. Volcano plots enriched sgRNAs in 25 mM fumaric acid condition. Black dots represent significantly enriched sgRNAs **C**. Representation of genes with significantly upregulated sgRNAs **D**. Distribution of activation, interference, and deletion sgRNAs per condition **E**. Percent distribution of *QDR2* activation sgRNAs in each fumaric acid condition. All conditions were done in triplicate.

Across all three fumaric acid conditions, we identified 90, 52, and 38 significantly enriched sgRNAs at 25 mM, 30 mM and 42 mM fumaric acid, respectively, based on a p-value threshold of < 0.05, an FDR value of < 0.05, and a log2 fold-change of at least 1.5 (Fig. 3B). In total, the enriched sgRNAs correspond to 86 unique genes across the three conditions (Fig. 3C). The top 10 gene hits for each condition are listed in Table 2. A list of all significantly enriched sgRNA can be found in Table S8. A clear trend emerged in the distribution of enriched sgRNAs, with activation sgRNAs comprising the vast majority across all fumaric acid concentrations (Fig. 3D). Activation sgRNAs constituted 98.98 %, 99.92 %, and 99.98 % of enriched sgRNAs in the 25 mM, 30 mM, and 42 mM fumaric acid conditions, respectively, compared to 62.3 % in the 0 mM control (Fig. 3D). Among the enriched activation sgRNAs, guides targeting *QDR2* were highly overrepresented (Fig. 3E). Activation sgRNAs for *QDR2* account for 90 %, 95 % and 98 % of enriched sgRNAs in the 25 mM, 30 mM and 42 mM conditions, respectively (Fig. 3E). Of the six activation sgRNAs designed for *QDR2*, four were significantly upregulated at 25 mM (Fig. 3B, green stars). While most of the highly enriched guides are activation sgRNAs, a small number corresponded to deletions (Table S8). Specifically, in the 25 mM condition, 81 of the significantly enriched sgRNAs were activation sgRNAs, 9 targeted gene deletion, and none were observed for interference.

**Table 2.** Top 10 list of sgRNAs significantly enriched in 25 mM, 30 mM and 42 mM fumaric acid growth conditions. LFC = Log2 fold change.

| Rank | 25mM | LFC | 30mM | LFC | 42mM | LFC |
| --- | --- | --- | --- | --- | --- | --- |
| 1 | actQDR2 | 10.604 | actQDR2 | 10.831 | actQDR2 | 10.972 |
| 2 | actQDR2 | 10.564 | actQDR2 | 10.269 | actQDR2 | 9.7333 |
| 3 | actFAS1 | 6.6233 | actUBP14 | 6.0777 | actUBP14 | 5.5647 |
| 4 | actUBP14 | 6.5932 | actFAS1 | 5.3817 | actIMD4 | 4.9176 |
| 5 | actQDR2 | 6.4336 | actIMD4 | 5.2305 | actCSE1 | 4.5154 |
| 6 | actIMD4 | 5.9393 | actQDR2 | 4.785 | actYPR1 | 3.7074 |
| 7 | actYPR1 | 5.1933 | actYPR1 | 4.0725 | actLYS21 | 3.5609 |
| 8 | delATP7 | 4.4575 | actCSE1 | 4.0243 | actFAS1 | 3.5103 |
| 9 | actACO2a | 4.3621 | actLYS21 | 3.8844 | actKLMA_20588 | 3.0591 |
| 10 | actQDR3 | 4.3215 | actRPN1 | 3.5368 | actQDR2 | 3.0521 |

### 3.4. Validation of Hits from CRISPR-AID Enrichment

To validate the results of the CRISPR-AID library selection, we individually tested the top 10 candidate genes with enriched sgRNAs under fumaric acid stress from each library. This resulted in a list of 12 unique gene perturbations, of which 11 were activation, and one was a deletion. To mimic activation in a CRISPR-Cas free system, the corresponding target genes were overexpressed from a plasmid using a strong promoter. For the single deletion target *ATP7* was deleted in the Y-1190Δ*FUM1* strain using CRISPR-Cas9 and then cured of the CRISPR-Cas9 plasmid. Growth rate analysis revealed that, while most candidate genes generated no significant improvement in fumaric acid tolerance, overexpression of the DHA1-family multidrug efflux pumps *QDR2* and *QDR3,* as well as deletion of ATP7, the stator arm of the F_1_F_0_ ATPase, exhibited notably higher growth rates at 20 mM fumaric acid compared to their respective control strains (Fig. 4A,B). At 20 mM and 42 mM fumaric acid, both *QDR2* and *QDR3* overexpression resulted in increased growth over EV, however, with an overall lower growth rate in 42 mM. *QDR2* overexpression caused an 8.7-fold increase in the growth rate at 42 mM fumaric acid compared to EV, reaching 0.09 h^-1^. A similar pattern was observed with *QDR3,* which had a growth rate of 0.14 h^-1^ compared to 0.01 h^-1^ for the EV (Fig. 4A). Due to the greater fumaric acid tolerance conferred by *QDR2* and *QDR3* overexpression compared to *ATP7* deletion, we focused our efforts on these transporters. We tested a *QDR2* and *QDR3* overexpression strain to determine if co-expression of both QDR genes could promote growth closer to the solubility limit of fumaric acid. We integrated *QDR2* and *QDR3* at known safe harbour sites (Rajkumar et al., 2019) and tested the growth of the strains in SC-Lactose supplemented with fumaric acid. The highest growth rates were measured in the *QDR2* integrated strain and in the *QDR2* integrated strain with one copy of *QDR3* (Fig. 4C). In both cases where *QDR3* was integrated into both alleles, maximum specific growth rate decreased in the absence of fumaric acid, but growth rate was still significantly higher than the Δ*FUM1* mutant in the presence of 42 mM fumaric acid (Fig. 4C). To further explore the role of these two genes in fumaric acid tolerance, growth rates were measured from Δ*QDR2* and Δ*QDR3* mutants of Y-1190Δ*FUM1* (Fig. 4D). Deletion of *QDR2*, whether alone or in combination with *QDR3*, significantly decreased the growth rate in 15 mM fumaric acid from 0.07 h^-1^ to 0.02 h^-1^ (Fig. 4D), while deletion of *QDR3* alone did not significantly change the growth rate under these conditions (Fig. 4D).

**Figure 4.**
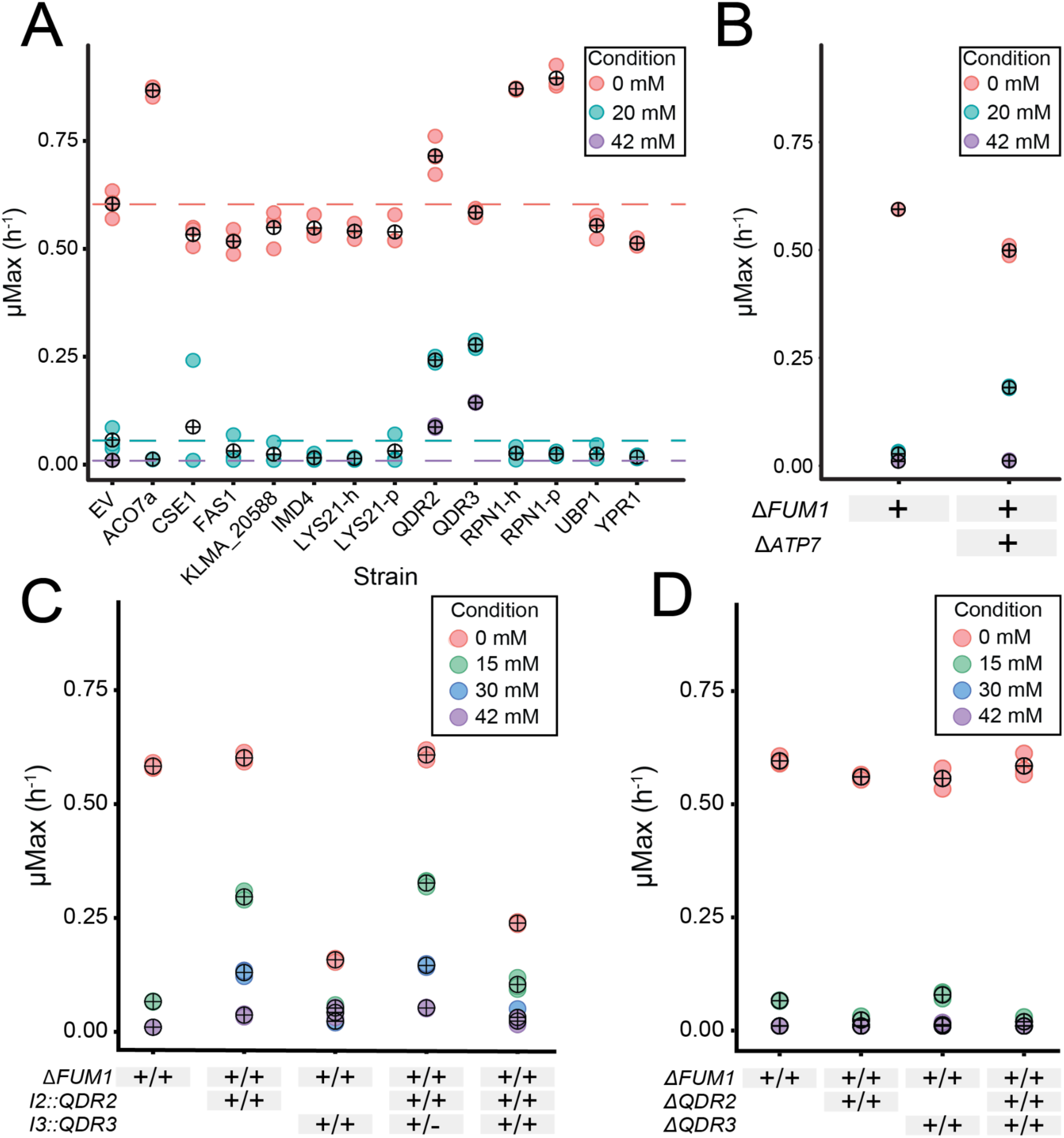
Validation of CRISPR-AID library hits in Y-1190Δ*FUM1* strain. Growth rates of the top hits from the CRISPR-AID screen in fumaric acid for **A.** activation and **B**. deletion. **C**. Growth rates of the integrated and **D.** deletion strains for *QDR2* and *QDR3*. Each coloured circle represents a data point, the black circles with crosses represent the mean of those data points.

### 3.5. Fumaric Acid Production

One of the main goals of this work was to create a fumaric acid production strain that is resistant to high concentrations of fumaric acid. Due to their role as multidrug efflux pumps and H+ antiporters in *S. cerevisiae* (Gbelska et al., 2006), we hypothesized that in *K. marxianus QDR2* and *QDR3* function as organic acid efflux pumps and that their overexpression would lead to higher fumaric acid production. To test this, we grew our overexpression strains in both SC-Lactose and 0.5x supplemented permeate, a lactose-rich feedstock, and measured metabolite profiles. Our results demonstrated that overexpression of *QDR2* in strain Y-1190Δ*FUM1* significantly increased fumaric acid production from 0.05 ± 0.00 g L^-1^ OD_600_^-1^ to 0.32 ±0.01 g L^-1^OD_600_^-1^, while *QDR3* overexpression resulted in slightly lower specific titers, reaching 0.13 ±0.02 g L^-1^ OD_600_^-1^ when grown in SC-Lactose media (Fig 5A). When grown in 0.5x permeate, we also saw significant increases in fumaric acid titers from 0.05 ±0.01 g L^-1^ OD_600_^-1^ to 0.26 ±0.02 g L^-1^OD_600_^-1^ and 0.21 ±0.00 g L^-1^ OD_600_^-1^ for *QDR2* and *QDR3*, respectively (Fig 5A).

**Figure 5.**
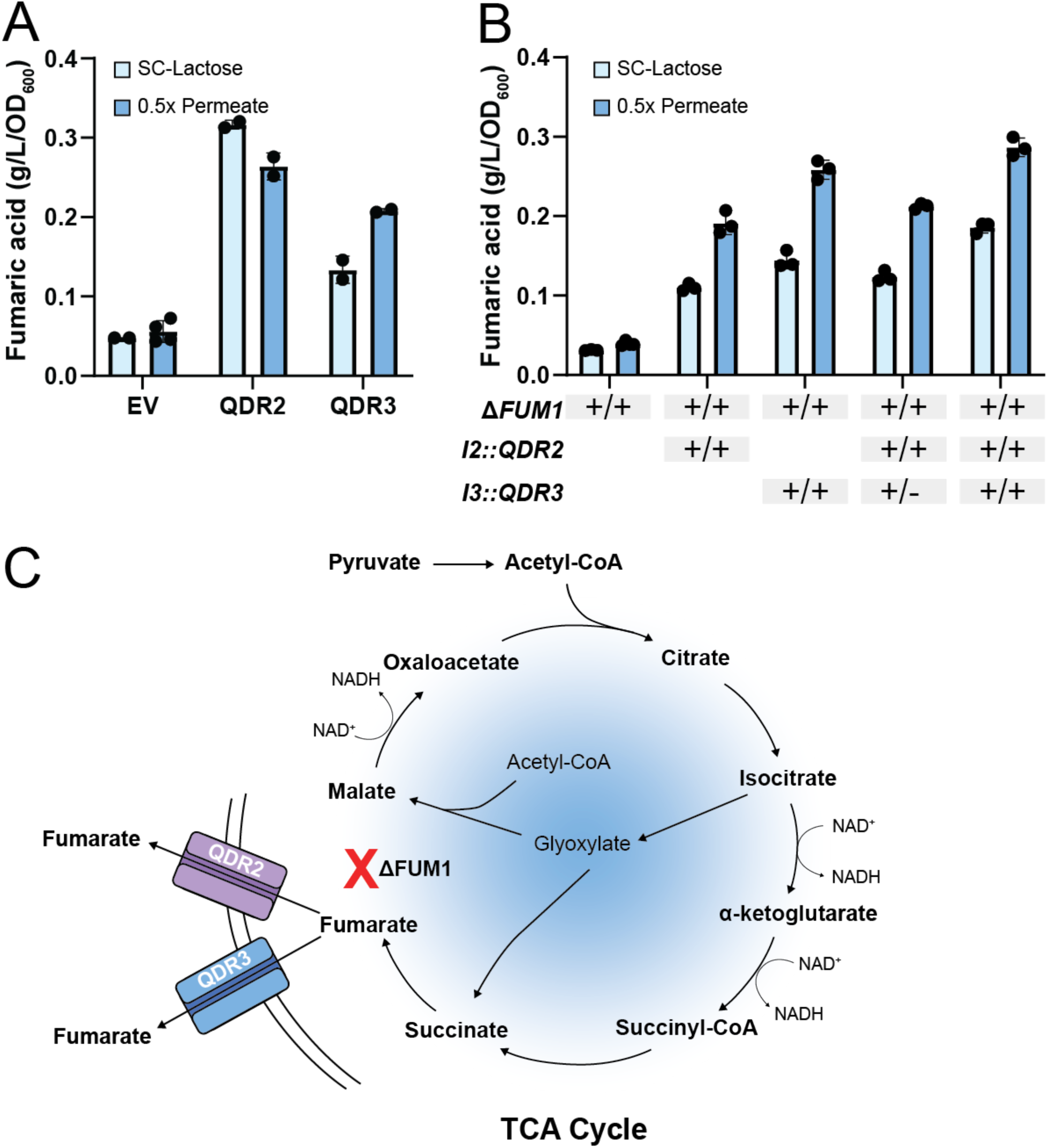
Effect of *QDR2* and *QDR3* on fumaric acid production in Y-1190Δ*FUM1* **A**. Peak fumaric acid specific titers for *QDR2* and *QDR3* over expressed on plasmid **B**. Fumaric acid specific titers at 48 h with *QDR2* and/or *QDR3* integrated into I2 and I3 safe harbour sites. **C**. Schematic of increased fumaric acid production in *K. marxianus* with overexpression of *QDR2* and *QDR3*. Fumaric acid was analysis on an HPLC were in duplicate (A) or triplicate (B) and normalized to OD_600_ measurements. This strain of *K. marxianus* is diploid. The + symbols represent how many copies were introduced into the genome. Error bars denote standard deviation.

To confirm these findings in a plasmid-free system, we repeated the fumaric acid production experiment in our *QDR2* and *QDR3* integrated strains. The genomic integration of *QDR2* and *QDR3* both under the strong promoter pNC1 resulted in titers of 0.19 ±0.00 g L^-1^ OD_600_^-1^ and 0.26 ±0.01 g L^-1^ OD_600_^-1^, respectively when grown in 0.5x supplemented permeate (Fig. 5B). When overexpressed together, we observed titers of 0.21±0.00 g L^-1^ OD_600_^-1^ for a single copy of *QDR3* and 0.29±0.01 g L^-1^ OD_600_^-1^ for the homozygous integration (Fig. 5B). The trends were the same when these strains were grown in SC-Lactose, but the overall titers were lower with *QDR2* and *QDR3* co-overexpression producing 0.19 ±0.01 g L^-1^ OD_600_^-1^.

While titers were the highest for the homozygous integration of *QDR2* and *QDR3* when normalized to biomass, the *QDR3* integrated strains produced less overall fumaric acid (Fig. S3). This is due to the overall lower growth of *QDR3* integrated strains (Fig. 4C). The highest titer we achieved was in the *QDR2* and single-copy *QDR3* integrated strain at 0.89±0.003 g L^−1^, which is ∼7.7 mM, much lower than 42 mM fumaric acid concentration we selected for (Fig. S3, Fig 3A). At 0 mM fumaric acid we observe a severe growth defect in *QDR3* integrated strains (Fig. 4C), which suggests that while *QDR3* overexpression may enhance fumaric acid production and tolerance at high concentrations, it may also impose a burden that limits cell growth in the absence of product accumulation.

To evaluate the impact of the Δ*FUM1, I2::QDR2, I3::QDR3* overexpression strain under controlled growth conditions, we scaled production up from 125 mL Erlenmeyer shake flasks to a 3 L fed-batch bioreactor. The *QDR2* and *QDR3* co-overexpression strain exhibited a striking improvement in fumaric acid production, reaching a maximum titer of 2.16 g L^-1^ at 89 hr, an 8.3-fold increase over the control Y-1190Δ*FUM1* strain at 0.26 g L^-1^ under identical conditions (Fig. 6) Both strains produced substantial amounts of acetic acid, with maximum titers of 8.14 g L^-1^ in the Δ*FUM1* and 12.39 g L^-1^ in the Δ*FUM1, I2::QDR2, I3::QDR3* overexpression strain (Fig. 6). Following the onset of exponential feeding at 49 h, neither strain exhibited long-term efficient lactose utilization, as evidenced by lactose accumulation commencing at ∼72 h in the Δ*FUM1* strain (Fig. 6A) and ∼90 h in the QDR overexpression strain (Fig. 6B). This result, together with the elevated acetic acid levels, suggests a potential metabolic bottleneck arising from the deletion of *FUM1*.

**Figure 6.**
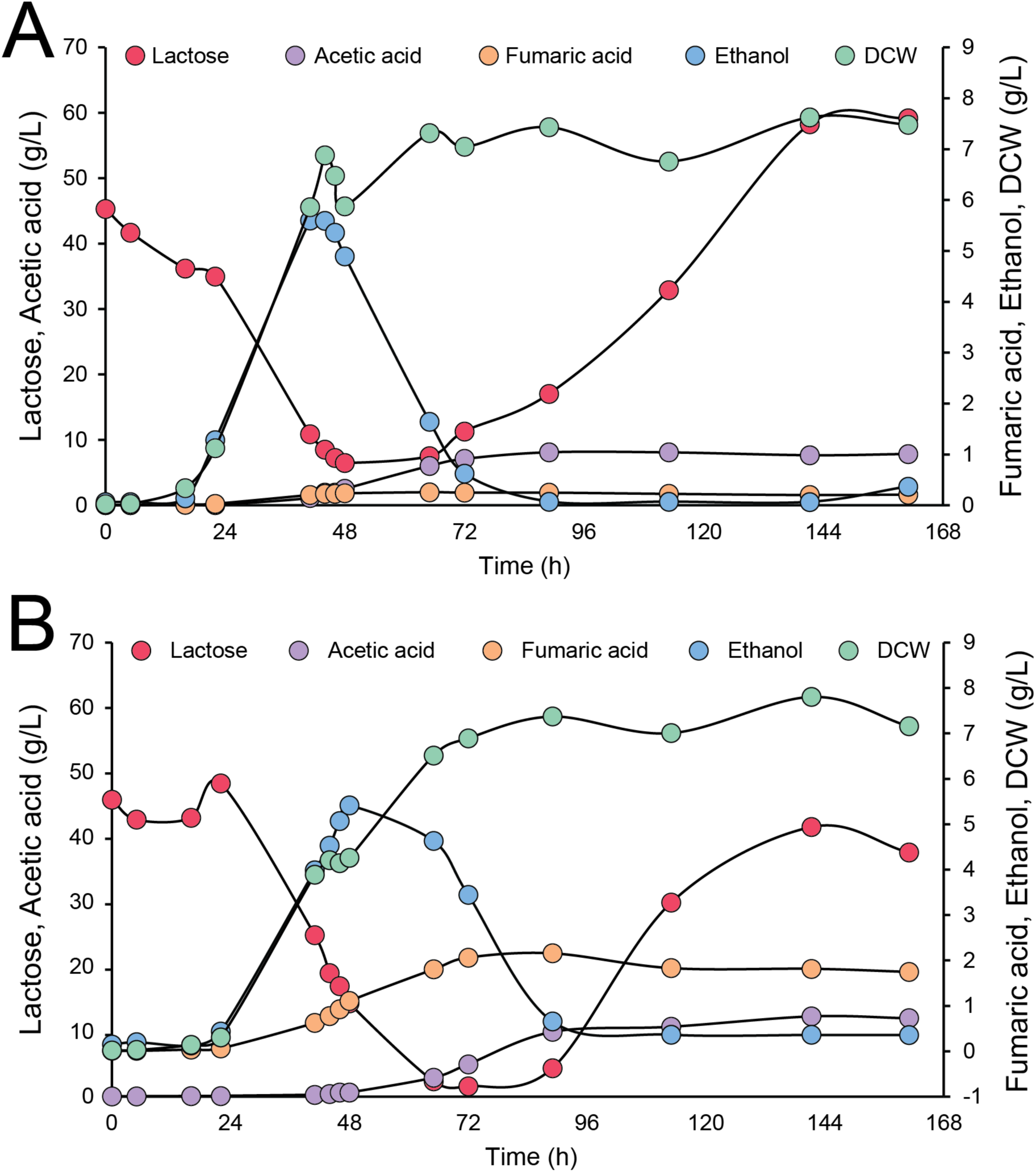
Growth and metabolite profiles of *K. marxianus* Y-1190 **A.** Δ*FUM1* and **B.** Δ*FUM1*, I2::*QDR2*, I3::*QDR3*. Fed-batch fermentation in supplemented permeate with pH maintained at 4. Fermentation profiles of dissolved oxygen, feed pump, and pH can be found in Figure S5.

## 4. Discussion

In this study, we investigated *K. marxianus* for fumaric acid production using lactose-rich dairy permeate as a sustainable feedstock. Deletion of the fumarate hydratase *FUM1* increased fumaric acid titers but compromised acid tolerance (Fig. 1). To address this, we applied a tri-functional CRISPR-AID library to unbiasedly identify genetic perturbations that improve acid tolerance. Through this approach, we identified activation of *QDR2* and *QDR3* gene expression, and deletion of *ATP7*, as beneficial modifications. Notably, overexpression of *QDR2* and *QDR3* also enhanced fumaric acid production to 2.16 g L^-1^, 8.3-fold over the *FUM1* deletion alone when grown in a fed-batch bioreactor (Fig. 6).

Much like in *S. cerevisiae* and *E. coli*, deletion of *FUM1* in *K. marxianus* increased fumaric acid accumulation (Song et al., 2013; G. Xu et al., 2012b) (Fig. 1), however, this modification also led to a reduction in growth in the presence of fumaric acid stress, a phenotype not previously reported. Notably, Xu *et al*. predicted that a *FUM1* deletion might reduce growth rate in *S. cerevisiae* based on a flux balance analysis, but experimentally observed no significant difference between the wild type and *FUM1*-deleted strains (G. Xu et al., 2012b). *FUM1* deletion represents only a preliminary step in the metabolic engineering of yeast for fumaric acid production. On its own, this perturbation resulted in modest titers of 0.610 ± 0.031 g L^-1^ in *S. cerevisiae* (G. Xu et al., 2012b). Many *S. cerevisiae* studies have implemented more comprehensive strategies, including transporter overexpression, rerouting flux toward fumaric acid, downregulation of competing pathways such as ethanol and glycerol production, and DNA scaffolds to spatially organize enzymes in close proximity (Chen et al., 2016; G. Xu et al., 2012a; Xu et al., 2013). These efforts have yielded fumaric acid titers ranging from 3.18 g L^-1^ to 33.13 g L^-1^ (Chen et al., 2016; G. Xu et al., 2012a; Xu et al., 2013). In *K. marxianus*, the reduced fumaric acid tolerance caused by the Δ*FUM1* deletion made enhancing tolerance a necessary prerequisite before advanced engineering strategies could be effectively implemented.

Unlike classical strain engineering approaches, CRISPR-based screens offer traceable interrogation of multiple genetic perturbations within one system. In addition, these functional approaches are especially powerful in non-conventional yeasts where much of the genome annotations are unknown or inferred from distant homology. While many CRISPR-based tools have been developed for *K. marxianus* (Bever et al., 2022; Cernak et al., 2018; Lee et al., 2018; Löbs et al., 2018; Nambu-Nishida et al., 2017; Rajkumar et al., 2019; Thornbury et al., 2025), to date, only one genome-wide knockout library has been constructed in *K. marxianus* (Robertson et al., 2025). To determine CRISPR-AID functionality in *K. marxianus*, we targeted mNG for activation, interference and deletion and demonstrated all three expected gene expression perturbations (Fig. 2A). Increased (2.6-to 4.6-fold) and decreased mNG fluorescence (2.9-to 5.7-fold) fell within the range reported in *S. cerevisiae*, with previous studies reporting 2.3-to 5-fold increases and 3-to 10-fold decreases in gene expression for activation and interference, respectively (Jensen et al., 2017). Similarly in the non-conventional yeast *Pichia pastoris*, CRISPRa and CRISPRi have been shown to yield 3.5-fold increases and 3.3-fold decreases, respectively (Liao et al., 2021). Prior CRISPRi studies in *K. marxianus* reported knockdown efficiencies between 1.5- and 5.6-fold (Löbs et al., 2018). Together, these results demonstrate that our CRISPR-AID system achieves gene modulation in *K. marxianus* at levels comparable to those reported in other yeast systems.

CRISPR-AID was originally applied in *S. cerevisiae* to enhance furfural tolerance in a genome-wide screen (Lian et al., 2019). In that study, most enriched guides corresponded to gene interference, with activation guides as the second most common and only a single deletion guide highlighted per condition. In contrast, our screen for fumaric acid tolerance revealed a strong enrichment for activation guides, comprising over ∼99% of hits at all fumaric acid concentrations, compared to 62% in the 0 mM control (Fig. 3D). Lian *et al*. identified hits that included interference of a negative regulator of the transporter *PDR5, PDR1* (Nishida-Aoki et al., 2015). In our case, top hits included activation of the multidrug transporters *QDR2* and *QDR3* (Fig. 4) whereas activation of *PDR5,* emerged in our screen at positions #30, #15, and #13 in the 25, 30, and 42 mM conditions, respectively (Table S8).

In our study, *QDR2* activation sgRNA accounted for 98% of all activation guide reads (Fig. 3) and was enriched ∼350-fold over the next most abundant validated hit, a *QDR3* activation sgRNA (Fig. 4). A previous application of genome-wide CRISPR-AID screening also found that activation guides were consistently enriched across three iterative screening rounds (Dong et al., 2021). For positive selection pooled library screens, it is common to have a narrow set of enriched hits. In an *E. coli* expression library screen for ethanol tolerance, three dominant hits were identified as being more abundant than the rest (Woodruff et al., 2013), and in a *S. cerevisiae* CRISPR deletion screen for furfural tolerance, one primary guide was strongly enriched along with two additional, less abundant hits (Bao et al., 2018). While this type of enrichment is typically desired, particularly under strong selection, it can also lead to population bottlenecks where a single dominant perturbation outcompetes others, potentially obscuring additional functionally relevant variants (Bock et al., 2022). This dynamic likely contributed to our own findings, in which we validated only two additional hits outside of the dominant enrichment for *QDR2* activation (Fig. 4A). It is possible that the fumaric acid concentrations applied for selection were overly stringent, leading to early bottlenecking of the library and overrepresentation of *QDR2* sgRNA. However, the CRISPR-AID study that relied solely on a fluorescent biosensor without applying direct selection pressure, still exhibited strong enrichment for a single dominant activation guide (Dong et al., 2021). This suggests that factors beyond selection stringency may drive the emergence of dominant hits, even in the absence of overt selective constraints.

*QDR2* and *QDR3* are genes encoding putative multidrug transporters previously characterized in *S. cerevisiae* as exporters involved in detoxification and efflux (Tenreiro et al., 2005; Vargas et al., 2006). Their activation in *K. marxianus* improved acid tolerance up to the solubility limit of fumaric acid, but also enhanced fumaric acid production, suggesting that these transporters may reduce intracellular toxicity of fumaric acid through export (Fig. 4, Fig. 5C). *QDR2* and *QDR3* are part of the Major Facilitator Superfamily (MFS), specifically, they are 12-spanner H+ antiporters in the *DHA1* family (Goffeau et al., 1997). Despite their similar names, *QDR2* and *QDR3* are from different clusters of the DHA1 family and share only 23.4 % identity at the amino acid level, the majority of the matching sequence being across the transmembrane domains (Dias et al., 2010) (Fig. 7). In *S. cerevisiae*, *QDR2* functions as a promiscuous cation/H⁺ antiporter with roles in potassium uptake (Vargas et al., 2006), tolerance to cation stresses (Ríos et al., 2013), and export of amino acids (Kapetanakis et al., 2021). Its broad substrate range, together with our finding that overexpression enhances growth in fumaric acid (Fig. 4AC) while deletion impairs it (Fig. 4D), suggests that *QDR2* may also export the anionic fumarate. QDR3 similarly confers multi-drug resistance (Tenreiro et al., 2005), exports threonine (Kapetanakis et al., 2021), and is implicated in tolerance to several dicarboxylic acids (Pereira et al., 2019), though paradoxically its deletion can improve acetic acid tolerance (Ma et al., 2015). The distinct yet overlapping substrate profiles and stress responses of QDR2 and QDR3 likely underly the enhanced fumaric acid tolerance we observed when both were overexpressed.

**Figure 7.**
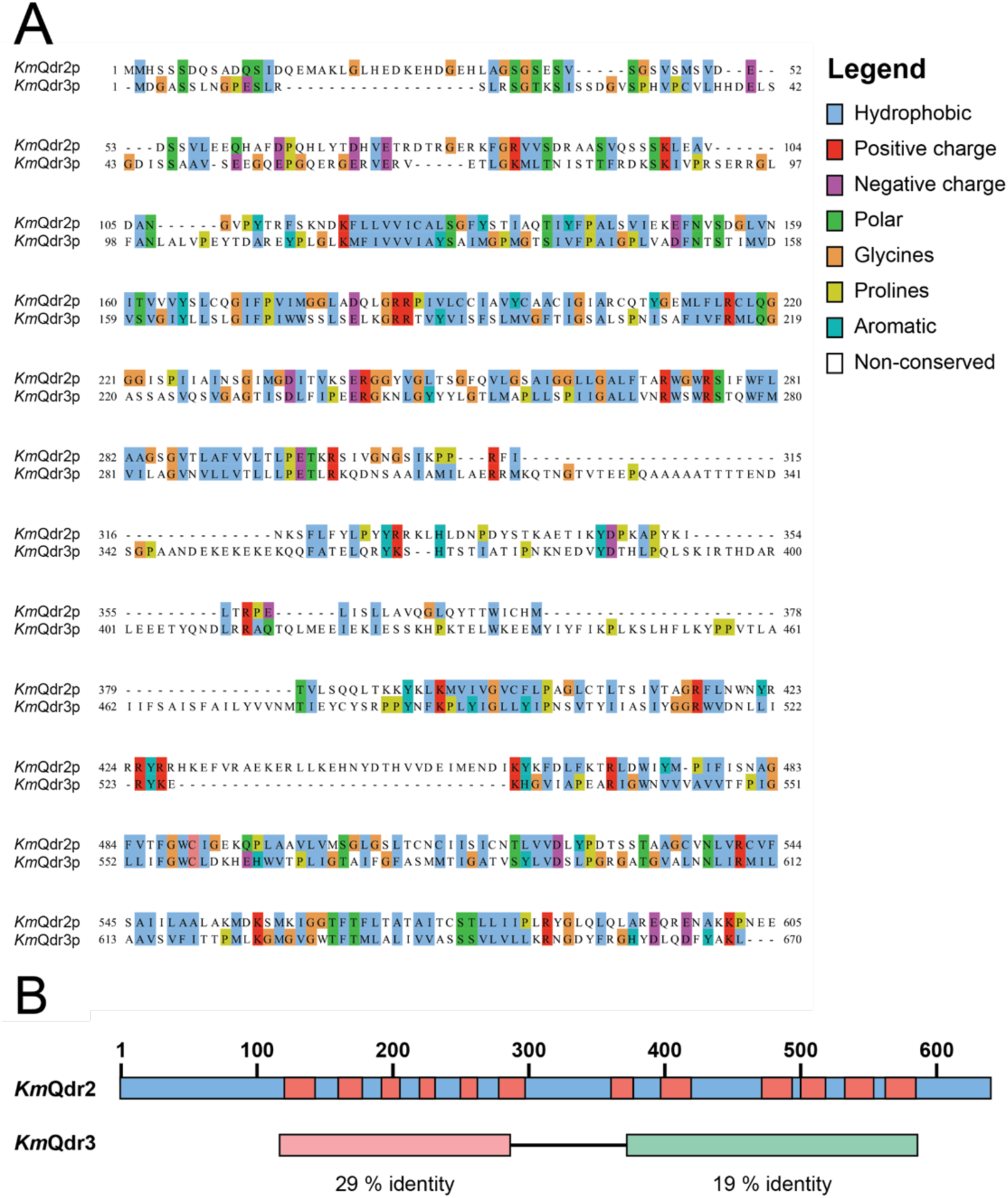
Alignment of *K. marxianus* Qdr2p and Qdr3p. **A**. Protein alignment at amino-acid level. **B**. A summary alignment with transmembrane domains highlighted in red, and areas of highest identity highlighted in pink and green. In *S. cerevisiae*, acid efflux is a common response to acid stress, typically mediated by members of the ABC and MFS transporter superfamilies (Sá-Correia and Godinho, 2022). Key ABC transporters include *SNQ2* for decanoic acid (Legras et al., 2010), and *PDR12* for sorbic, propionic, benzoic, octanoic, and levulinic acids (Holyoak et al., 1999; Legras et al., 2010; Nygård et al., 2014). Among MFS transporters, several DHA1 and DHA2 members, including *AQR1*, *TPO1-4*, and *AZR1* export a range of short-to medium-chain acids (Godinho et al., 2017; Legras et al., 2010; Tenreiro et al., 2002, n.d.; Zhang et al., 2022). Within this framework, *QDR2* and *QDR3*, members of the DHA1 subfamily of MFS transporters, align well with other characterized acid efflux transporters, supporting their roles in mediating organic acid tolerance.

While deletion mutants leading to enhanced cellular function are less common than leading to reduced cellular function, examples such as deletion of *YBR238C* or *FOB1* in *S. cerevisiae* enhance yeast lifespan through enhanced mitochondrial function or replication of rDNA, respectively (Alfatah et al., 2024; Defossez et al., 1999; Kaeberlein et al., 2005). In our case, the deletion of *ATP7*, which encodes a subunit of the stator arm of the mitochondrial F₁F₀ ATPase (Norais et al., 1991), emerged as an unexpected but significant hit in our screen, conferring increased tolerance to 20 mM fumaric acid (Fig. 4B). The F₁F₀ ATPase is the mitochondrial ATPase required for ATP generation through oxidative phosphorylation (Weber and Senior, 1997). Disruption of ATPase complexes in general, including ATP7, are not classically associated with increasing acid tolerance. The identification of ATP7 deletion in our study underscores the strength of an unbiased, genome-wide approach in revealing non-obvious genetic determinants of stress tolerance.

We observe high titers of fumaric acid in *QDR3* integrated strains when normalized to biomass (Fig. 5), but overall lower growth and final titers (Fig. S3). When grown in 0.5x permeate, but not SC media, our fumaric acid producing strains also produce high titers of acetic acid, reaching 5.6 g L^-1^ at maximum (Fig. S4), possibly decreasing the fitness of the more susceptible *QDR3* overexpressed integrants. This effect may be linked to differences in nitrogen sources: while both media contain ∼ 30 g L^-1^ lactose, SC media uses ammonium sulphate for the nitrogen source, whereas 0.5x supplemented permeate uses urea (Thornbury et al., 2025). Urea metabolism catabolism releases CO_2_ which can dissolve into the cytosol to increase the pH, rather than acidifying it, allowing for further accumulation of acetic acid before cell death (Hensing et al., 1995; Yang et al., 2021). Supporting this idea, the combined *QDR2* and *QDR3* overexpression strain produced acetic acid at 12.39 g L^-1^ when grown in a bioreactor with the pH controlled to 4.0, likely mitigating some acid stress (Fig. 6).

The high concentrations of acetic acid observed in both shake flask and bioreactor experiments (Fig. S4, Fig. 6) indicate a metabolic imbalance, likely caused by the Δ*FUM1* mutation disrupting the TCA cycle (Fig. 1). Acetic acid accumulation may result from excess acetyl-CoA production or from overflow metabolism to generate additional NADH (Cui et al., 2017; Pentjuss et al., 2017), and the reduced TCA flux due to *FUM1* deletion likely contributes to both mechanisms. While *QDR2* and *QDR*3 overexpression enhances fumaric acid production, these transporters cannot fully mitigate the toxicity caused by accumulating acetic acid, highlighting a remaining metabolic limitation in the engineered strains.

With fumaric acid-tolerant *K. marxianus* strain in hand, we are now well-positioned to pursue targeted metabolic engineering strategies to further improve fumaric acid production. The identification and integration of fumaric acid tolerance-enhancing genes such as *QDR2* and *QDR3* provide a robust chassis for production. Future work will focus on reducing overflow metabolism and fine-tuning cofactor balance to maximize fumarate yield and reduce acetic acid. Together, these efforts will enable the construction of a high-performing production strain and expand the utility of *K. marxianus* as a platform for organic acid biosynthesis from industrial waste streams.

## Acknowledgements

We thank Dr. Aida Tafrishi and Rory Coles who helped with troubleshooting the CHOPCHOP package and downstream analysis on python.

## Supplementary Material

1. M_Thornbury_SupplementaryMaterial.docx includes supplementary figures 1-5 and supplementary tables 1-4, 6-8. Due to the size of the table, Table S5 has been provided in the following excel document
2. M_Thornbury_Table S5.xlsx Supplementary Table 5 that includes all sgRNA included in the genome-wide CRISPR-AID libraries.

## Author Contributions

Conceptualization of the project was carried out by VJJM, MW, MT, and AK, while formal analysis was performed by MT and AK. Funding acquisition was managed by VJJM, and MW, and investigation was conducted by MT, RO, LK, AK, and RA. The project administration was facilitated by VJJM, MW, MT, and AK. Resources and supervision were provided by VJJM, and MW. Validation was conducted by MT, AK, RO, and LK, and visualization was done by MT. The original draft of the writing was produced by MT, with the review and editing process carried out by VJJM, MW, MT, AK, and RO.

## Declaration of generative AI and AI-assisted technologies in the writing process

During the preparation of this work the authors used ChatGPT to check grammar and improve clarity. After using this tool, the authors reviewed and edited the content as needed and take full responsibility for the content of the published article.

