## Supplemental Figures and Tables for "Tri-Functional CRISPR Screen Reveals Overexpression of *QDR2* and *QDR3* Transporters Increase Fumaric Acid Production in *Kluyveromyces marxianus*"

**Supplementary Figures**

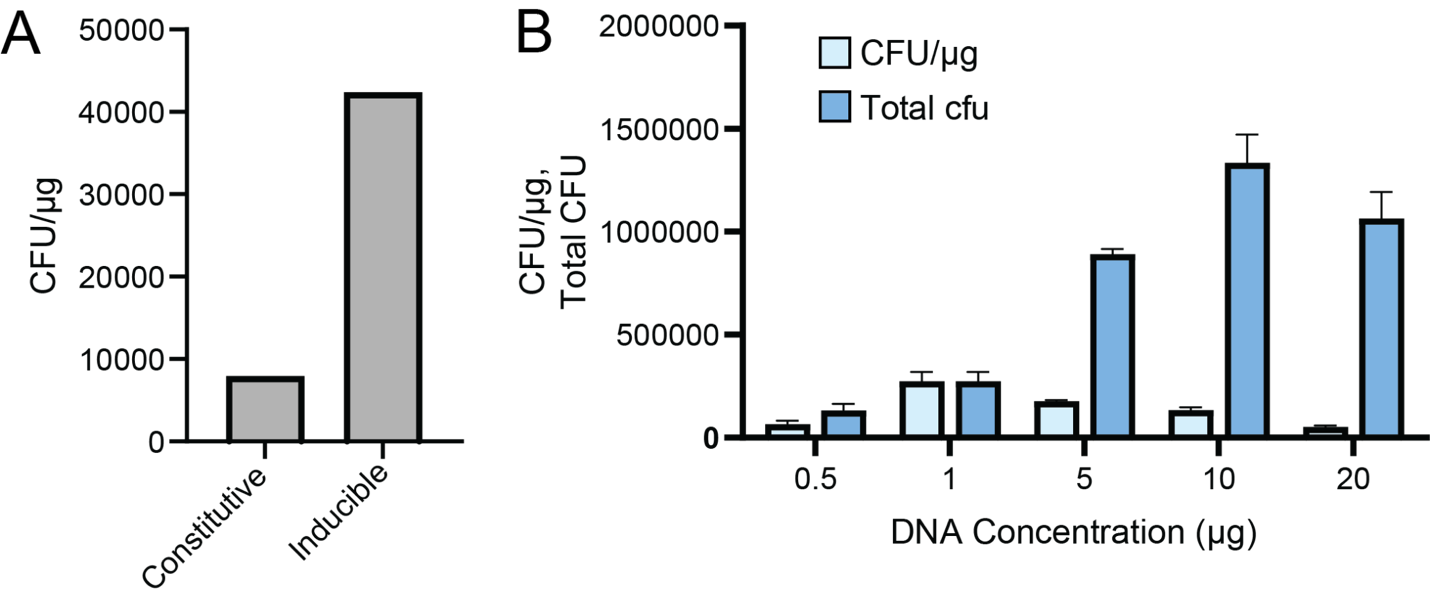

**Figure S1**. Transformation efficiency with CRISPR-AID system **A**. Comparison of transformation efficiency of *K. marxianus* Y-1190 with the tri functional Cas cassette expressed constitutively (left), or under inducible promoters (right) **B**. Transformation efficiency in the inducible Cas system across multiple DNA concentrations. Error bars represent the standard deviation between two biological replicates.

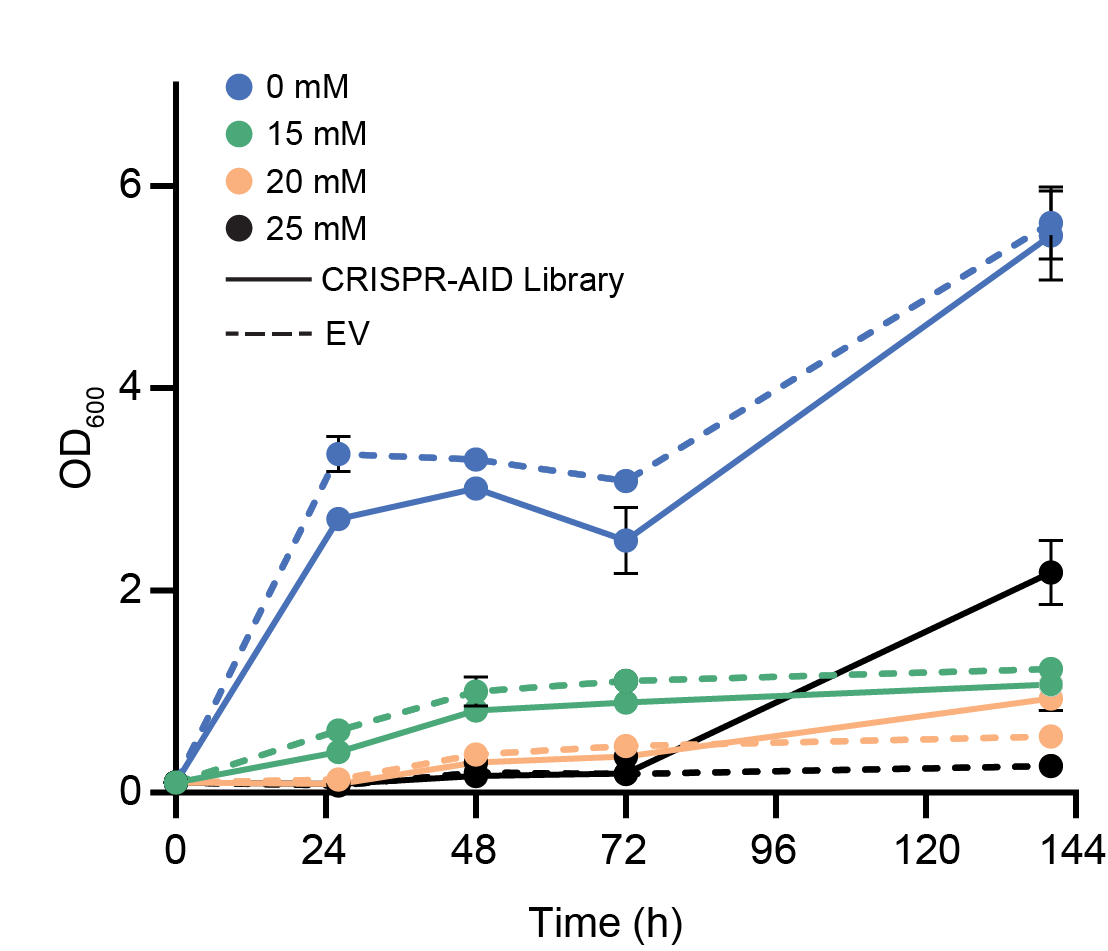

**Figure S2**. Testing fumaric acid concentrations to be used in CRISPR-AID enrichment experiments. In triplicate, MTK074 with an EV plasmid, or with the transformed genome-wide CRISPR-AID sgRNA library was grown in the presence of multiple concentrations of fumaric acid over six days. Each dot is the mean of three biological replicates and error bars represent standard deviation.

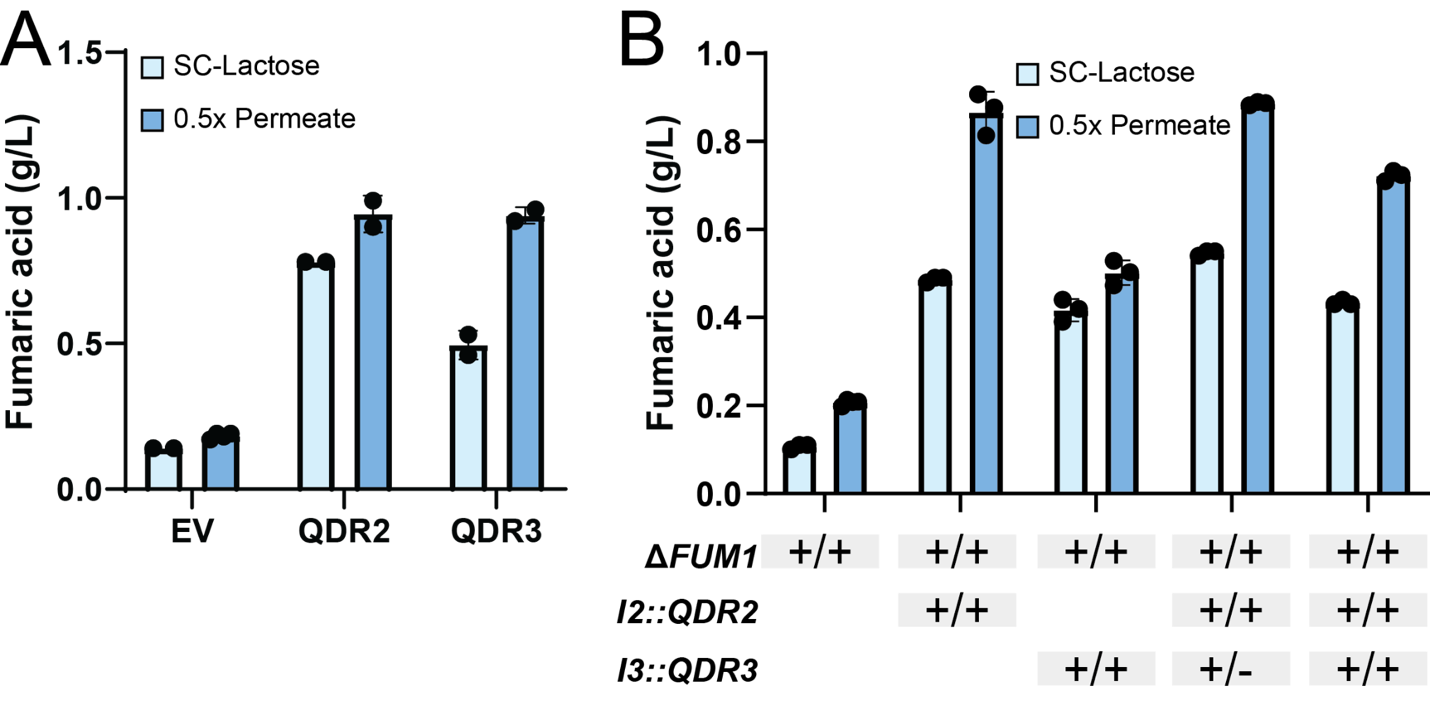

**Figure S3**. Fumaric acid titers in SC-Lactose and 0.5x Permeate with QDR2 and QDR3 overexpressed **A**. on plasmid or **B**. by integration. Fumaric acid was measured on an HPLC using an Aminex 87-H column (Bio-Rad) in duplicate (A) or triplicate (B). This strain of *K. marxianus* is diploid. The + symbols represent how many copies were introduced into the genome. Error bars denote standard deviation.

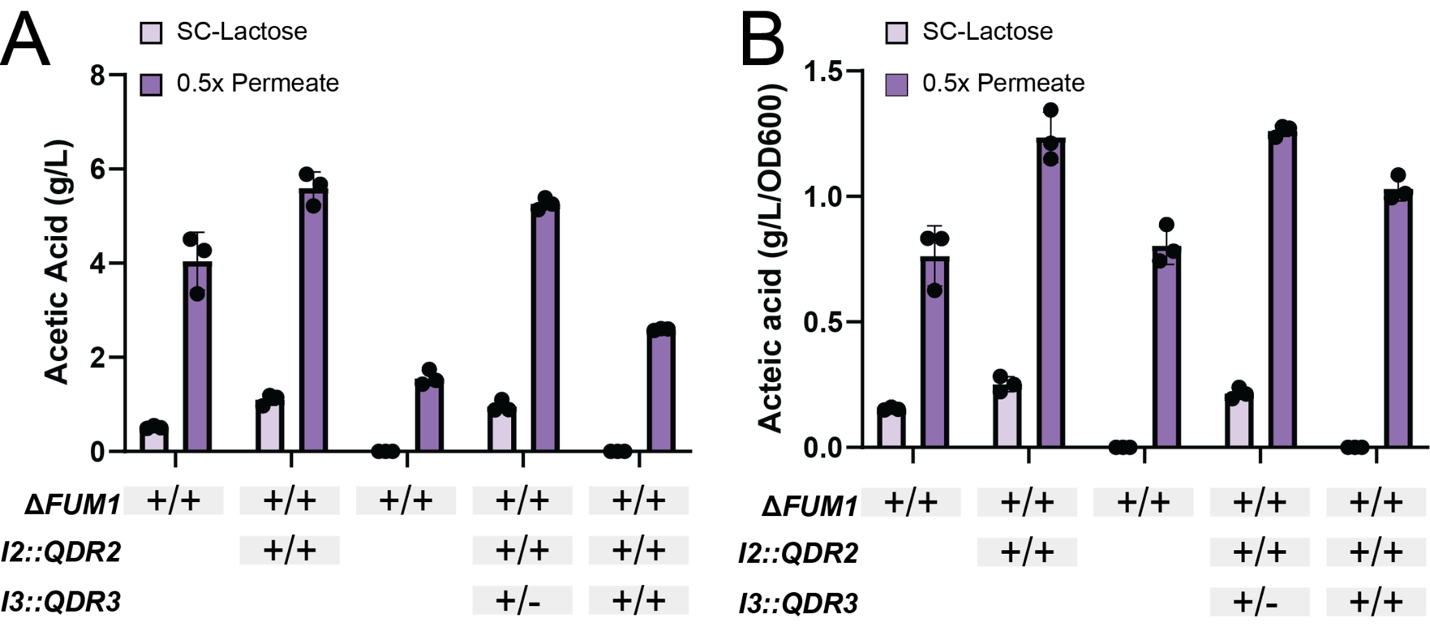

**Figure S4**. Acetic acid **A**. Total titers and **B**. Specific production normalized to OD_600._ Acetic acid was measured on an HPLC using an Aminex 87-H column (Bio-Rad) in triplicate. This strain of *K. marxianus* is diploid. The + symbols represent how many copies were introduced into the genome. Error bars denote standard deviation.

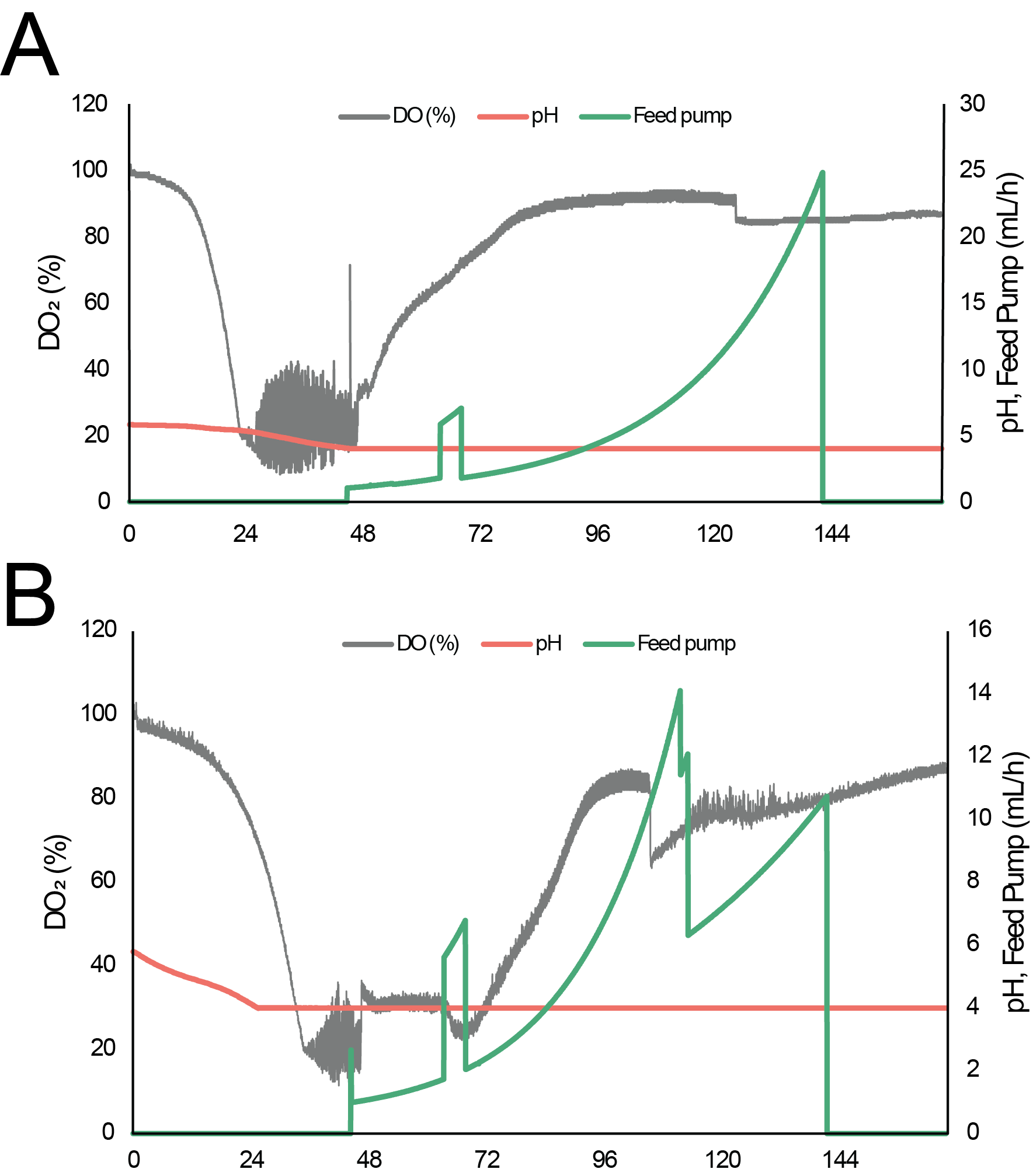

**Figure S5.** Fed-batch fermentation profiles of *K. marxianus* Y-1190 **A.** Δ*FUM1* and **B.** *ΔFUM1*, I2::*QDR2*, I3::*QDR3*. Bioreactor was set up in batch mode until lactose was depleted at ~49 hr, at which time fed-batch was started using 200 g L^-1^ lactose. pH was maintained at 4. Process data were recorded and analyzed using BioXpert V2 (Applikon).

**Supplementary Tables**

**Table S1.** List of plasmids used in this study

| **Plasmid ID** | **Parental Plasmid** | **Characteristic** | **Selection** | **Reference** | **Addgene** |
| --- | --- | --- | --- | --- | --- |
| Kmk.C1 | - | Minimal ARS and centromeric sequence | Cam | Rajkumar, 2019 | 125059 |
| Kmk.I2L | - | Integration left arm downstream of KmBDH1 and KmBDH2 | Cam | Rajkumar, 2019 | 125031 |
| Kmk.I2R | - | Integration right arm downstream of KmBDH1 and KmBDH2 | Cam | Rajkumar, 2019 | 125064 |
| Kmk.I3L | - | Integration left arm between KmSWF1 and KmARO1 | Cam | Rajkumar, 2019 | 125032 |
| Kmk.I3R | - | Integration right arm between of KmSWF1 and KmARO1 | Cam | Rajkumar, 2019 | 125065 |
| Kmk.I4L | - | Integration left arm downstream of KmHSP104 | Cam | Rajkumar, 2019 | 125033 |
| Kmk.I4R | - | Integration right arm downstream of KmHSP104 | Cam | Rajkumar, 2019 | 125066 |
| Kmk.P2 | - | CBS 6556 PDC1 promoter | Cam | Rajkumar, 2019 | 125035 |
| Kmk.T3 | - | CBS 6556 KMXK_A03020 terminator | Cam | Rajkumar, 2019 | 125055 |
| Kmk.T4 | - | CBS 6556 PDC1 terminator | Cam | Rajkumar, 2019 | 125056 |
| Kmk.T5 | - | CBS 6556 PGK1 terminator | Cam | Rajkumar, 2019 | 125057 |
| p426*-Lb | - | Scaffold for LbCas12a for S. cerevisiae | Amp, -URA | Lian et al., 2019 | 136992 |
| p426*-Sa | - | Scaffold for SaCas9 for S. cerevisiae | Amp, -URA | Lian et al., 2019 | 136994 |
| p426*-Sp | - | Scaffold for SpCas9 for S. cerevisiae | Amp, -URA | Lian et al., 2019 | 136993 |
| pAID6 | - | CRISPR-AID Integration casette for S. cerevisiae | Amp, G418 | Lian et al., 2019 | 136989 |
| pAK007 | pYTK090 | I2 GFP dropout landing pad | Kan | This work | - |
| pAK012 | pMTKM01 | SpCas9-sgRNA1-sgRNA2-double break in I2 landing pad | Amp, Nat | This work | - |
| pAK034 | pYTK090 | I4 Landing Pad | Kan | This work | - |
| pAK104 | pMT189 | Inducible CRISPR-AID Integration Casette: KmTEF3tetO2prom-dLbCas12-VP-ScADH1term, KmTDH3tetO2prom-SaCas9-KmPDC1term, and KmPGK1tetO2prom-dSpCas9-RD1152-KmA3020term | Kan, G418 | This work | - |
| pBB316 | - | mScarlet CDS | Cam | Derivative of [Bean et al., 2022](https://academic.oup.com/g3journal/article/12/8/jkac154/6609175) | - |
| pJD005 | pMTKM01 | URA3 double guide set | Amp, Nat | This work | - |
| pJet1.2 | - | CloneJET PCR Cloning Kit | Amp | Thermofisher, K1231 | - |
| pMT124 | YTK083 | Scaffold for SpCas9 in a level1 plasmid | Amp, -URA | This work | - |
| pMT149 | pMT172 | Sg4_mNeon_a guide in scaffold | Amp, -URA | This work | - |
| pMT150 | pMT172 | Sg5_mNeon_a guide in scaffold | Amp, -URA | This work | - |
| pMT151 | pMT172 | Sg6_mNeon_a guide in scaffold | Amp, -URA | This work | - |
| pMT152 | pMT177 | Sg7_mNEON guide in scaffold | Amp, -URA | This work | - |
| pMT153 | pMT177 | Sg8_mNeon guide in scaffold | Amp, -URA | This work | - |
| pMT154 | pMT124 | Sg1_mNeon_i guide in scaffold | Amp, -URA | This work | - |
| pMT155 | pMT124 | Sg2_mNeon_i guide in scaffold | Amp, -URA | This work | - |
| pMT156 | pMT124 | Sg3_mNeon_i guide in scaffold | Amp, -URA | This work | - |
| pMT172 | YTK083 | Scaffold for LbCas12 in a level1 plasmid, with GFP dropout | Amp, -URA | This work | - |
| pMT177 | YTK083 | Scaffold for SaCas9 in a level1 plasmid, with GFP dropout | Amp, -URA | This work | - |
| pMT188 | pJet1.2 | URA large::mNeon | Amp | This work | - |
| pMT189 | - | I3 landing pad with selection | Kan, G418 | This work | - |
| pMT193 | pMTKM01 | sgRNA plasmid targeting I3 locus | Amp, Nat | This work | - |
| pMT195 | pMT189 | Repair template to integrate the CRISPR-AID cassette: KmPGK1p-dLbCas12-VP-ScADH1t, KmTDH3p-SaCas9-KmPDC1t, and KmTEF3p-dSpCas9-RD1152-KmA3020t | Kan, G418 | This work | - |
| pMT230 | - | Destination plasmid for multiplexed guide and scaffold combinations. SNR52 promoter and SUP4 terminator | Amp, -URA | This work | - |
| pMT238 | pYTK090 | I3 Landing Pad without selection | Kan | This work | - |
| pMT249 | pMT230 | Combo CRISPR Test 1 (A) | Amp, -URA | This work | - |
| pMT250 | pMT230 | Combo CRISPR Test 2 (B) | Amp, -URA | This work | - |
| pMT251 | pMT230 | Combo CRISPR Test 3 (C) | Amp, -URA | This work | - |
| pMT252 | pMT230 | PDC1 activation with Sg4 act guide | Amp, -URA | This work | - |
| pMT253 | pMT230 | PDC1 inhibition with Sg2 Int guide | Amp, -URA | This work | - |
| pMT254 | pMT230 | mNeon Green deletion with Sg7 del guide | Amp, -URA | This work | - |
| pMT266 | pAK034 | I4 Donor: mTurquoise | Kan | This work | - |
| pMT267 | pMT230 | CRISPR-AID EV | Amp, -URA | This work | - |
| pMT273 | pMT230 | SOD1p guide act A | Amp, -URA | This work | - |
| pMT274 | pMT230 | SOD1p guide act B | Amp, -URA | This work | - |
| pMT275 | pMT230 | SOD1p guide act C | Amp, -URA | This work | - |
| pMT276 | pMT230 | TEF3p guide int A | Amp, -URA | This work | - |
| pMT277 | pMT230 | TEF3p guide int B | Amp, -URA | This work | - |
| pMT278 | pMT230 | TEF3p guide int C | Amp, -URA | This work | - |
| pMT321 | YTK077 | golden gate G418 destination plasmid | amp,G418 | This work | - |
| pMT351 | - | Destination plasmid for guide and scaffold combinations. SNR52 promoter and SUP4 terminator | Amp, -URA | This work | - |
| pMT418 | pMTKM01 | QDR2 CDS guide set 1 | Amp, Nat | This work | - |
| pMT421 | pMT321 | pNC1_QDR2_ScADH1t | Amp, G418 | This work | - |
| pMT422 | pMT321 | pNC1_LYS21_primary_ScADH1t | Amp, G418 | This work | - |
| pMT423 | pMT321 | pNC1_LYS21_haplotig_ScADH1t | Amp, G418 | This work | - |
| pMT424 | pMT321 | pNC1_IMD4_ScADH1t | Amp, G418 | This work | - |
| pMT425 | pMT321 | pNC1_g3201_primary_ScADH1t | Amp, G418 | This work | - |
| pMT426 | pMT321 | pNC1_g3201_haplotig_ScADH1t | Amp, G418 | This work | - |
| pMT427 | pMT321 | pNC1_g2114_ScADH1t | Amp, G418 | This work | - |
| pMT428 | pMT321 | pNC1_CSE1_ScADH1t | Amp, G418 | This work | - |
| pMT429 | pMT321 | pNC1_FAS1_ScADH1t | Amp, G418 | This work | - |
| pMT430 | pMT321 | pNC1_UBP14_ScADH1t | Amp, G418 | This work | - |
| pMT432 | pAK007 | NC1-QDR2-ScADH1 I2 donor | Kan | This work | - |
| pMT433 | pMT321 | pNC1_QDR3_ScADH1t | Amp, G418 | This work | - |
| pMT434 | pMT321 | spacer Empty vector (EV) | Amp, G418 | This work | - |
| pMT435 | pMT321 | pNC1-YPR1-SCADH1t | Amp, G418 | This work | - |
| pMT443 | pMT238 | I3 Donor: pNC1-QDR3-PGKt | Kan | This work | - |
| pMT444 | pMTKM01 | QDR3 CDS guide set 1 | Amp, Nat | This work | - |
| pMT466 | pMT321 | pNC1-ACO7a-ScADH1t | Amp, G418 | This work | - |
| pMT467 | pMT321 | pNC1-RPN1-ScADH1t | Amp, G418 | This work | - |
| pMT468 | pMTKM01 | ATP7 guide 1 | Amp, NAT | This work | - |
| pMTKM01 | - | pCAS-Tyr-[gRNA: BsaI GFP dropout] with pan(OPT)ARS  replication origin | Amp, Nat | Thornbury et al., 2025 | 233473 |
| pMTKM24 | - | Level 0 plasmid MoClo with KM promoter PGK1 | Cam | Thornbury et al, 2025 | 233496 |
| pMTKM25 | - | Level 0 plasmid MoClo with KM promoter TEF3 | Cam | Thornbury et al, 2025 | 233497 |
| pMTKM26 | - | Level 0 plasmid MoClo with KM promoter TDH3 | Cam | Thornbury et al, 2025 | 233498 |
| pMTKM28 | YTK001 | Y-1190 NC1 promoter | Cam | Thornbury et al, 2025 | 233500 |
| pMTKM30 | - | TDH3-tetO2p level 0 Moclo part 2 (TetR repressed, with 2XtetO sequences) | Cam | Thornbury et al, 2025 | 233502 |
| pMTKM31 | - | TEF3-tetO2p level 0 Moclo part 2 (TetR repressed, with 2XtetO sequences) | Cam | Thornbury et al, 2025 | 233503 |
| pMTKM32 | - | PGK1-tetO2p level 0 Moclo part 2 (TetR repressed, with 2XtetO sequences) | Cam | Thornbury et al, 2025 | 233504 |
| pYTK001 | - | Part Plasmid Entry Vector | Cam | Lee et al., 2015 | 65108 |
| pYTK032 | - | mTurquoise2 CDS | Cam | Lee et al., 2015 | 65139 |
| pYTK053 | - | ScADH1 terminator | Cam | Lee et al., 2015 | 65160 |
| pYTK077 | - | Encodes KanamycinR | Cam, G418 | Lee et al., 2015 | 65184 |
| pYTK083 | - | Encodes AmpR-ColE1 | Amp | Lee et al., 2015 | 65190 |
| pYTK090 | - | Encodes KanR-ColE1 | Kan | Lee et al., 2015 | 65197 |

**Table S2.** List of oligonucleotides used in this study

| **Application** | **OligoID** | **Nucleotide sequence (5’ – 3’)** |
| --- | --- | --- |
| Amplify dSpCas9-RD1152 from pAID6 | MT174 | GCATCGTCTCATCGGTCTCATATGatggacgtcccaaagaag |
| Amplify dSpCas9-RD1152 from pAID6 | MT175 | ATGCCGTCTCAGGTCTCAGGATttacgcaactggaacagatg |
| Amplify SaCas9 from pAID6 | MT176 | GCATCGTCTCATCGGTCTCATATGatggggaaacggaactac |
| Amplify SaCas9 from pAID6 | MT177 | ATGCCGTCTCAGGTCTCAGGATTTAcgagactttcctcttcttcttg |
| Amplify dLbCas12a-VP from pAID6 | MT178 | GCATCGTCTCATCGGTCTCATATGccaccatggctcctcc |
| Amplify dLbCas12a-VP from pAID6 | MT179 | ATGCCGTCTCAGGTCTCAGGATtcacagcaaggctgagaaatc |
| SaCas9 diagnostic primer | MT191 | tctctttgccggttgttc |
| SaCas9 diagnostic primer | MT192 | aagatctgctgaataatccattc |
| dLbCas12a-VP diagnostic primer | MT193 | gtggaatgccgagtatgac |
| dLbCas12a-VP diagnostic primer | MT194 | gcaagctgtatatgttccag |
| dLbCas12a-VP diagnostic primer | MT195 | gtatcagaagttcgagaagatg |
| dLbCas12a-VP diagnostic primer | MT196 | cattggacgattttgatctgg |
| dSpCas9-RD1152 diagnostic primer | MT201 | ttcagcttcaatgattaagcg |
| dSpCas9-RD1152 diagnostic primer | MT202 | gttcgacgataaagtgatgaag |
| dSpCas9-RD1152 diagnostic primer | MT203 | atatctgaatgccgtggtg |
| dSpCas9-RD1152 diagnostic primer | MT204 | caagtacttcgacaccacc |
| Amplifies 900 bp upstream of URA3 | MT293 | GCAAGGGAATTTGCCACC |
| Amplifies 900 bp upstream of URA3, 30 bp homology to prPDC1 | MT294 | tccattattcacgctgtatattcgctggatCCTCCTTTGATTAGTTTATGGGC |
| Amplifies 900 bp downstream of URA3, 30 bp homology to PDC1term | MT297 | caagttacaagttacaagttacaagttaccAGAGTTCTCCGAGAACAAGC |
| Amplifies 900 bp downstream of URA3 | MT298 | TGGTATGCTTCTTGTGCATTC |
| Amplifies NeonGreen transcriptional unit with PDC1 prom&ter | MT295 | atccagcgaatatacagcg |
| Amplifies NeonGreen transcriptional unit with PDC1 prom&ter | MT296 | Tggtaacttgtaacttgtaacttg |
| mNeonGreen diagnostic primer | MT303 | atcttcggttccattaacgg |
| mNeonGreen diagnostic primer | MT304 | cgttaatggaaccgaagatg |
| short guide for inhibition of Δ URA::mNEON by CCTOP | MT324 | TGATCGAGTGTCTGGATCAGTTTTG |
| short guide for inhibition of Δ URA::mNEON by CCTOP | MT325 | AAACCAAAACTGATCCAGACACTCG |
| short guide for inhibition of Δ URA::mNEON by CCTOP | MT326 | TGATCATAATACAAATATACATTAA |
| short guide for inhibition of Δ URA::mNEON by CCTOP | MT327 | AAACTTAATGTATATTTGTATTATG |
| short guide for inhibition of Δ URA::mNEON by CCTOP | MT328 | TGATCAATACAAATATACATTAAAG |
| short guide for inhibition of Δ URA::mNEON by CCTOP | MT329 | AAACCTTTAATGTATATTTGTATTG |
| short guide for activation of Δ URA::mNEON by CCTOP | MT330 | AGATGAATACTGAATATCGAATAT |
| short guide for activation of Δ URA::mNEON by CCTOP | MT331 | AAAAATATTCGATATTCAGTATTC |
| short guide for activation of Δ URA::mNEON by CCTOP | MT332 | AGATTATATAAGTGGAGTGTCTGG |
| short guide for activation of Δ URA::mNEON by CCTOP | MT333 | AAAACCAGACACTCCACTTATATA |
| short guide for activation of Δ URA::mNEON by CCTOP | MT334 | AGATATTTTATTCTGGTTCCTGCG |
| short guide for activation of Δ URA::mNEON by CCTOP | MT335 | AAAACGCAGGAACCAGAATAAAAT |
| short guide for deletion of Δ URA::mNEON by CCTOP | MT336 | TGATCTCAAGGGCGAAGCCCAAGTT |
| short guide for deletion of Δ URA::mNEON by CCTOP | MT337 | AAACAACTTGGGCTTCGCCCTTGAG |
| short guide for deletion of Δ URA::mNEON by CCTOP | MT338 | TGATCGAACCGAAGATGTGCAACTC |
| short guide for deletion of Δ URA::mNEON by CCTOP | MT339 | AAACGAGTTGCACATCTTCGGTTCG |
| pSOD1 activation guide_1 | MT557 | taccactaaaccacttgcgcATGGAGGAAGGCAATTGAAGAAGatctacactta |
| pSOD1 activation guide_2 | MT558 | taccactaaaccacttgcgcGAAAAGGAGCTGAAAATGACTTAatctacacttagtagaaatt |
| pSOD1 activation guide_3 | MT559 | taccactaaaccacttgcgcAGGCGTAGCGCAACCCCTGTGGAatctacacttagtagaaatt |
| Universal forward primer for dLbCas12a guides | MT556 | ttcgattccgggcttgcgcaaatttctactaagtgtagat |
| Universal reverse guide for dSpCas9 guides | MT560 | taccactaaaccacttgcgcaaaagcaccgactcggtgccactttttcaagttgataacggactagccttattttaacttg |
| pTEF1 interference guide_1 | MT561 | ttcgattccgggcttgcgcaTCAATTTTGAATAATTAGAGgttttagagctagaaatagcaagttaaaataaggctagtc |
| pTEF1 interference guide_2 | MT562 | ttcgattccgggcttgcgcaTTTTTCCTTCTGTTTAGTACgttttagagctagaaatagcaagttaaaataaggctagtc |
| pTEF1 interference guide_3 | MT563 | ttcgattccgggcttgcgcaAAAATTTACATTATCGAAAAgttttagagctagaaatagcaagttaaaataaggctagtc |
| Universal reverse guide for SaCas9 guides | MT573 | taccactaaaccacttgcgcatcaaaaaaatctcgccaacaagttgac |
| mNeonGreen deletion guide_1 | MT565 | ttcgattccgggcttgcgcaTCAAGGGCGAAGCCCAAG |
| mNeonGreen deletion guide_2 | MT566 | ttcgattccgggcttgcgcaGAACCGAAGATGTGCAAC |
| Amplification of sgRNA library with heterogeneity spacers, universal reverse | MT788 | TCGTCGGCAGCGTCagatgtgtataagagacagtgatatgccatatatgcagtggtg |
| Amplification of sgRNA library with heterogeneity spacers, NGS1 | MT789 | GTCTCGTGGGCTCGGagatgtgtataagagacagCTAtctccgcagtgaaagataaatgatc |
| Amplification of sgRNA library with heterogeneity spacers, NGS2 | MT790 | GTCTCGTGGGCTCGGagatgtgtataagagacagTGACtctccgcagtgaaagataaatgatc |
| Amplification of sgRNA library with heterogeneity spacers, NGS3 | MT791 | GTCTCGTGGGCTCGGagatgtgtataagagacagGTCTAtctccgcagtgaaagataaatgatc |
| Amplification of sgRNA library with heterogeneity spacers, NGS4 | MT792 | GTCTCGTGGGCTCGGagatgtgtataagagacagACTGTGtctccgcagtgaaagataaatgatc |
| Amplification of sgRNA library with heterogeneity spacers, NGS5 | MT793 | GTCTCGTGGGCTCGGagatgtgtataagagacagTAGTCAGtctccgcagtgaaagataaatgatc |
| Amplification of sgRNA library with heterogeneity spacers, NGS6 | MT794 | GTCTCGTGGGCTCGGagatgtgtataagagacagTGCtctccgcagtgaaagataaatgatc |
| Amplification of sgRNA library with heterogeneity spacers, NGS7 | MT795 | GTCTCGTGGGCTCGGagatgtgtataagagacagCTGAtctccgcagtgaaagataaatgatc |
| Amplification of sgRNA library with heterogeneity spacers, NGS8 | MT796 | GTCTCGTGGGCTCGGagatgtgtataagagacagCTGAAtctccgcagtgaaagataaatgatc |
| Amplification of sgRNA library with heterogeneity spacers, NGS9 | MT797 | GTCTCGTGGGCTCGGagatgtgtataagagacagATCGACtctccgcagtgaaagataaatgatc |
| Amplification of sgRNA library with heterogeneity spacers, NGS10 | MT798 | GTCTCGTGGGCTCGGagatgtgtataagagacagTAGTTGCtctccgcagtgaaagataaatgatc |
| Amplification of QDR2 (g4399) from K.marxianus Y-1190 | MT812 | GCATCGTCTCATCGGTCTCATatgatgcatagtagtagtgatc |
| Amplification of QDR2 (g4399) from K.marxianus Y-1190 | MT813 | ATGCCGTCTCAGGTCTCAGGATctatagagctgagaaatttgattgc |
| Amplification of double guide set for QDR2 | MT820 | AAAAGGTCTCAGACTTTACTCTGACCCTGACCCCGCCgttttagagctagaaatagcaagttaaaataag |
| Amplification of double guide set for QDR2 | MT821 | TACGGGTCTCTAAACttggcatccactgcttctagtgcgcaagcccggaatc |
| QDR2 diagnostic primer | MT855 | atgatgcatagtagtagtgatcaaag |
| QDR2 diagnostic primer | MT825 | gccagacctcccataataac |
| Amplification of IMD4 from K. marxianus Y-1190 | MT835 | GCATCGTCTCATCGGTCTCATATGccaggtaagctattgg |
| Amplification of IMD4 from K. marxianus Y-1190 | MT836 | ATGCCGTCTCAGGTCTCAGGATttagttgtgtagacgtttttcg |
| Amplification of QDR3 from K. marxianus Y-1190 | MT837 | GCATGAAGACAATCGGTCTCATatggacggggcaagttc |
| Amplification of QDR3 from K. marxianus Y-1190 | MT858 | ATGCGAAGACAAGGTCTCAGGATttattcaagcttagcatagaaatcttg |
| Amplification of LYS21 from K. marxianus Y-1190 | MT839 | GCATCGTCTCATCGGTCTCATATGtctgtgaactcaaatccg |
| Amplification of LYS21 from K. marxianus Y-1190 | MT840 | ATGCCGTCTCAGGTCTCAGGATtcataactccgaatggaaatttttg |
| Amplification of g2114 from K. marxianus Y-1190 | MT841 | GCATCGTCTCATCGGTCTCATatggatctagaggcagatatctc |
| Amplification of g2114 from K. marxianus Y-1190 | MT842 | ATGCCGTCTCAGGTCTCAGGATctactcgatttgtttttgctttgg |
| Amplification of YPR1 from K. marxianus Y-1190 | MT843 | GCATCGTCTCATCGGGAAGACAATatgtcacatttgaagccatc |
| Amplification of YPR1 from K. marxianus Y-1190 | MT856 | ATGCCGTCTCAGGTCGAAGACAAGGATctattcaaatgtcttgaaagtacc |
| Amplification of CSE1 from K. marxianus Y-1190 | MT845 | ATCGCGTCTCATatggccgatttagagtcag |
| Amplification of CSE1 from K. marxianus Y-1190 | MT846 | ATCGCGTCTCAGGATctatcgagcacataattttgttaatg |
| Amplification of FAS1 from K. marxianus Y-1190 | MT847 | ATCGCGTCTCATatgagctcatctgcaagacc |
| Amplification of FAS1 from K. marxianus Y-1190 | MT848 | ATCGCGTCTCAGGATttattggttaacaatcttttcccag |
| Amplification of g3201 from K. marxianus Y-1190 | MT849 | GCATCGTCTCATCGGTCTCATatgaatccaacgagtgctc |
| Amplification of g3201 from K. marxianus Y-1190 | MT850 | ATGCCGTCTCAGGTCTCAGGATctagaccggactattatcaataaatttag |
| Removal of internal bsmbi sites in IMD4 | MT851 | ATTACGTCTCaggacggtaagcgtttgaag |
| Removal of internal bsmbi sites in IMD4 | MT852 | ATTACGTCTCagtccctgtagaaatattcacc |
| Amplification of UBP14 from K. marxianus Y-1190 | MT853 | ATCGCGTCTCATatgtctgactttgatatatgtgac |
| Amplification of UBP14 from K. marxianus Y-1190 | MT857 | ATCGCGTCTCAGGATtcattccaatagggtgaagaag |
| QDR3 diagnostic primers | MT877 | agaaagaatattggaacaatcactag |
| QDR3 diagnostic primers | MT878 | catgaaccattgtgtcgatc |
| Amplification of double guide set for ATP7 | MT892 | AAAAGGTCTCAGACTTTCTTACCGGTCAACTTCAAAGgttttagagctagaaatagcaagttaaaataag |
| Amplification of double guide set for ATP7 | MT893 | TACGGGTCTCTAAACctattgaggctttcgaatcgtgcgcaagcccggaatc |
| ATP7 diagnostic primer | MT896 | atgtctctagccaaatctgc |
| ATP7 diagnostic primer | MT897 | acataacggtcaaatcaccg |
| Amplification of RPN1 from K. marxianus Y-1190 | MT898 | GCATCGTCTCATCGGTCTCATatggctgaagataaagtgactaaag |
| Amplification of RPN1 from K. marxianus Y-1190 | MT899 | ATGCCGTCTCAGGTCTCAGGATttaattcacttcttctgtgtactttgg |
| Amplification of ACO7a from K. marxianus Y-1190 | MT900 | GCATCGTCTCATCGGTCTCATatgctagctgctagaagatc |
| Amplification of ACO7a from K. marxianus Y-1190 | MT901 | ATGCCGTCTCAGGTCTCAGGATtcatgtagatgagcggc |
| I2 diagnostic primers | AK831 | AGGTATATAAGCGAGAAATTGGC |
| I2 diagnostic primers | AK832 | TAATCACGGTGAACTGAATGC |
| I4 diagnostic primers | AK950 | tcctttcctgttcaatgcag |
| I4 diagnostic primers | AK951 | tgtgggtgtgtgtttttctc |
| I3 diagnostic primers | MT487 | CTTTACGTGATCGAGACCG |
| I3 diagnostic primers | MT488 | tctAGCCATTTGATATCAACAAAC |
| Blunt primers for first amplification of the AID library | MT736 | attccgggcttgcgca |
| Blunt primers for first amplification of the AID library | MT737 | taccactaaaccacttgcgc |
| Primers with overhangs for library insertion into entry vector | AK954 | GCATGGTCTCAAATCCAACGTTGCCATCGTTGGGCCCCCGGTTCGATTCCGGGCTTGCGCA |
| Primers with overhangs for library insertion into entry vector | AK961 | GCATGGTCTCACCCAACGATGGCAACGTTGGATTTTACCACTAAACCACTTGCGC |
| Amplification of double guide set for QDR3 | MT863 | AAAAGGTCTCAGACTTTGACGGGGCAAGTTCTTTAAAgttttagagctagaaatagcaagttaaaataag |
| Amplification of double guide set for QDR3 | MT864 | TACGGGTCTCTAAACCGCATCTGTATACTCCGGGAtgcgcaagcccggaatc |

**Table S3**. List of yeast strains used in this study

| **Species** | **Strain Name** | **Parental Strain** | **Genetic modification** | **Details** | **Selection** |
| --- | --- | --- | --- | --- | --- |
| K. marxianus | Y-1190 | - | - | Diploid yeast | - |
| K. marxianus | ROK002 | Y-1190 | ΔFUM1 |  | - |
| K. marxianus | MTK017 | Y-1190 | ΔURA3::mNeonGreen |  | - |
| K. marxianus | MTK019 | Y-1190 | ΔURA3::mNeongGreen, I3::CRISPR-AID | CRISPR-AID = KmPGK1p-dLbCas12-VP-ScADH1t, KmTDH3p-SaCas9-KmPDC1t, and KmTEF3p-dSpCas9-RD1152-KmA3020t | G418 |
| K. marxianus | MTK056 | MTK019 | ΔURA3::mNeonGreen, I3:: CRISPR-AID, I4::mturquoise (het) | CRISPR-AID = KmPGK1p-dLbCas12-VP-ScADH1t, KmTDH3p-SaCas9-KmPDC1t, and KmTEF3p-dSpCas9-RD1152-KmA3020t | G418 |
| K. marxianus | MTK063 | MTK056 | ΔURA3::mNeonGreen, I3:: CRISPR-AID, I4::mturquoise (het), I2::mScarlett , | CRISPR-AID = KmPGK1p-dLbCas12-VP-ScADH1t, KmTDH3p-SaCas9-KmPDC1t, and KmTEF3p-dSpCas9-RD1152-KmA3020t | G418 |
| K. marxianus | MTK072 | MTK017 | ΔURA3::mNeonGreen, I3::inducibleCRISPR-AID | inducibleCRISPR-AID=SSA3p-tetR-SV40-Inu1t-ConL1-TEF3tetO2p-LbCas12a-VP-ScADH1t-ConL2-TDH3tetO2p-SaCas9-PDC1t-ConL3-PGK1tetO2p-SpCas9-RD1152-KMX_A3020t-G418 | G418 |
| K. marxianus | MTK074 | MTK072 | ΔURA3::mNeonGreen, I3::inducibleCRISPR-AID, ΔFUM1 | inducibleCRISPR-AID=SSA3p-tetR-SV40-Inu1t-ConL1-TEF3tetO2p-LbCas12a-VP-ScADH1t-ConL2-TDH3tetO2p-SaCas9-PDC1t-ConL3-PGK1tetO2p-SpCas9-RD1152-KMX_A3020t-G418 | G418 |
| K. marxianus | MTK115 | ROK002 | ΔFUM1, ΔQDR2 |  | - |
| K. marxianus | MTK117 | ROK002 | ΔFUM1, I2:: pNC1-QDR2-ScADH1term |  | - |
| K. marxianus | MTK129 | ROK002 | ΔFUM1, I3::pNC1-QDR3-PGK1term |  | - |
| K. marxianus | MTK130het | MTK117 | ΔFUM1, I2:: pNC1-QDR2-ScADH1term, I3::pNC1-QDR3-PGK1term (het) |  | - |
| K. marxianus | MTK130ho | MTK117 | ΔFUM1, I2:: pNC1-QDR2-ScADH1term, I3::pNC1-QDR3-PGK1term (hom) |  | - |
| K. marxianus | MTK135 | MTK115 | ΔFUM1, ΔQDR2, ΔQDR3 |  | - |
| K. marxianus | MTK139 | ROK002 | ΔFUM1, ΔQDR3 |  | - |
| K. marxianus | MTK142 | ROK002 | ΔFUM1, ΔATP7 |  | - |

**Table S4.** sgRNA Scoring Rubric

| **Scoring** | **Activation** | **Interference** | **Deletion** |
| --- | --- | --- | --- |
| Efficiency (E) | (Kim et al., 2018) | (Xu et al., 2015) | (Doench et al., 2014) |
| Position (a) | a = \|x-250\|/250 | a =\|X-125\|/125 | If x/CDS <1/3, a=0  If 1/3 =<X/CDS<=2/3, a=0.2  If X/CDS>2/3, a =0.5 |
| GC Score (b) | if 40-60% b=0  if 30-40% or 60-70% b=0.2  if 20-30% or 70-80% b=0.4  if 10-20% or 80-90% b=0.6  if 0-10% or 90-100% b=0.8 | | |
| Off-target score (c) | c = (SM+MM0+MM1+MM2+MM3)/20 | | |
| PolyT & PolyG Score (d & e) | If consecutive T>4, d =1, else d=0  If consecutive G>5, e =1, else e =0 | | |
| Diversity Score (f) | If distance <10 bp, f=0  Else, f=1 | | |
| Total score (S) | S= (3+E-a-b-c)*d*e*f | | |

**Table S5.** Please see attached excel file.

**Table S6**. Genome-wide CRISPR-AID Library Coverage

|  | Library Size | *E. coli* transformants  (total cfu) | *E. coli* library coverage | *K. marxianus* Transformants (total cfu) | *K. marxianus* Library Coverage |
| --- | --- | --- | --- | --- | --- |
| Activation | 26185 | 4.84x10^5^ | 18.5 | 5.4x10^5^ | 20.6 |
| Interference | 26988 | 5.15x10^5^ | 19.1 | 8.9x10^5^ | 33.0 |
| Deletion | 18763 | 4.00x10^5^ | 21.3 | 3.2x10^5^ | 17.1 |
| Total | 71936 | 1.40x10^6^ | - | 1.75x10^6^ | - |

**Table S7**. Library Sequencing Read Coverage per Condition

| Condition | Replicate | Merged reads | Percent Assembled | Theoretical read coverage per guide |
| --- | --- | --- | --- | --- |
| 0 mM | A | 1080223 | 99.45 | 15.02 |
|  | B | 1145441 | 99.44 | 15.92 |
|  | C | 1027931 | 99.49 | 14.29 |
| 25 mM | A | 1288488 | 99.49 | 17.91 |
|  | B | 1248906 | 99.45 | 17.36 |
|  | C | 1187174 | 99.48 | 16.50 |
| 30 mM | A | 1115467 | 99.43 | 15.51 |
|  | B | 1300580 | 99.47 | 18.08 |
|  | C | 1229784 | 99.47 | 17.10 |
| 42 mM | A | 1062307 | 99.49 | 14.77 |
|  | B | 659048 | 99.37 | 9.16 |
|  | C | 899214 | 99.44 | 12.50 |
| Average |  | 1.10E+06 | 99.46 | 15.34 |

**Table S8.** Significantly enriched sgRNA from the genome-wide CRISPR-AID screen in fumaric acid.

| Condition | Gene name | sgRNA ID | Common Name | LFC | FDR |
| --- | --- | --- | --- | --- | --- |
| 25 mM Fumaric acid | g4399 | ACT21802 | QDR2 | 10.604 | 0 |
| 25 mM Fumaric acid | g4399 | ACT21806 | QDR2 | 10.564 | 0 |
| 25 mM Fumaric acid | g3577 | ACT16642 | FAS1 | 6.6233 | 0 |
| 25 mM Fumaric acid | g3178 | ACT14031 | UBP14 | 6.5932 | 0 |
| 25 mM Fumaric acid | g4399 | ACT21801 | QDR2 | 6.4336 | 0 |
| 25 mM Fumaric acid | g592 | ACT25286 | IMD4 | 5.9393 | 0 |
| 25 mM Fumaric acid | g2142 | ACT07423 | YPR1 | 5.1933 | 0 |
| 25 mM Fumaric acid | g4077 | DEL13335 | ATP7 | 4.4575 | 4.63E-209 |
| 25 mM Fumaric acid | g902 | ACT27254 | ACO2a | 4.3621 | 0 |
| 25 mM Fumaric acid | g547 | ACT24985 | QDR3 | 4.3215 | 0 |
| 25 mM Fumaric acid | g1501 | ACT03264 | LYS21 | 4.1276 | 0 |
| 25 mM Fumaric acid | g1123 | ACT00802 | NNK1 | 4.0638 | 4.71E-201 |
| 25 mM Fumaric acid | g292 | ACT12438 | KLMA_70317 | 4.0618 | 8.03E-282 |
| 25 mM Fumaric acid | g1640 | ACT04144 | RPN1 | 3.9984 | 0 |
| 25 mM Fumaric acid | g3201 | ACT14193 | KLMA_40132 | 3.9145 | 0 |
| 25 mM Fumaric acid | g3020 | ACT13106 | CSE1 | 3.8772 | 1.1226e-314 |
| 25 mM Fumaric acid | g3618 | ACT16917 | TOM1 | 3.8754 | 2.58E-268 |
| 25 mM Fumaric acid | g878 | ACT27085 | GIS2 | 3.8138 | 3.67E-154 |
| 25 mM Fumaric acid | g971 | ACT27711 | KLMA_10227 | 3.7693 | 5.21E-138 |
| 25 mM Fumaric acid | g2114 | ACT07232 | KLMA_20588 | 3.7518 | 0 |
| 25 mM Fumaric acid | g4278 | ACT21032 | GPI12 | 3.6949 | 9.50E-298 |
| 25 mM Fumaric acid | g2445 | ACT09339 | RRN3 | 3.6405 | 6.64E-167 |
| 25 mM Fumaric acid | g2043 | ACT06768 | PKP1 | 3.5818 | 2.60E-127 |
| 25 mM Fumaric acid | g1390 | DEL01718 | SNQ2 | 3.5325 | 5.70E-154 |
| 25 mM Fumaric acid | g1538 | ACT03500 | PEX32 | 3.5141 | 2.73E-196 |
| 25 mM Fumaric acid | g2642 | ACT10606 | CDA2 | 3.4765 | 4.36E-68 |
| 25 mM Fumaric acid | g267 | ACT10790 | KLMA_70292 | 3.4488 | 2.07E-71 |
| 25 mM Fumaric acid | g694 | ACT25965 | LAS21 | 3.4488 | 4.90E-153 |
| 25 mM Fumaric acid | g3308 | ACT14880 | PDR5 | 3.4192 | 3.08E-242 |
| 25 mM Fumaric acid | g4399 | ACT21805 | QDR2 | 3.4067 | 0 |
| 25 mM Fumaric acid | g659 | ACT25730 | GIP4 | 3.3736 | 1.02E-204 |
| 25 mM Fumaric acid | g4406 | ACT21855 | ZUO1 | 3.3221 | 1.14E-169 |
| 25 mM Fumaric acid | g3813 | ACT18126 | g3813 | 3.2926 | 1.67E-171 |
| 25 mM Fumaric acid | g2056 | ACT06851 | CAP2 | 3.285 | 5.19E-90 |
| 25 mM Fumaric acid | g2189 | ACT07724 | ARG81 | 3.279 | 3.12E-178 |
| 25 mM Fumaric acid | g1345 | ACT02252 | KLMA_10605 | 3.2759 | 4.94E-176 |
| 25 mM Fumaric acid | g3014 | ACT13062 | DUG1 | 3.2401 | 2.32E-148 |
| 25 mM Fumaric acid | g1291 | ACT01896 | HGT1 | 3.2201 | 1.21E-96 |
| 25 mM Fumaric acid | g2525 | ACT09854 | ATP16 | 3.1961 | 5.92E-187 |
| 25 mM Fumaric acid | g3700 | ACT17407 | VPS54 | 3.1134 | 5.24E-98 |
| 25 mM Fumaric acid | g407 | ACT19751 | PAC2 | 3.1113 | 1.06E-119 |
| 25 mM Fumaric acid | g547 | ACT24990 | QDR3 | 3.0754 | 1.87E-148 |
| 25 mM Fumaric acid | g2227 | ACT07985 | VMA2 | 3.019 | 3.84E-87 |
| 25 mM Fumaric acid | g1670 | ACT04334 | COX16 | 2.9979 | 2.77E-43 |
| 25 mM Fumaric acid | g4307 | ACT21221 | NSE1 | 2.9202 | 2.19E-83 |
| 25 mM Fumaric acid | g2521 | ACT09831 | RTG1 | 2.898 | 4.97E-177 |
| 25 mM Fumaric acid | g2989 | ACT12888 | SKI8 | 2.8409 | 8.11E-27 |
| 25 mM Fumaric acid | g3331 | ACT15031 | OCH1 | 2.8236 | 6.45E-68 |
| 25 mM Fumaric acid | g987 | ACT27810 | SDS23 | 2.7823 | 2.11E-52 |
| 25 mM Fumaric acid | g1362 | ACT02359 | YKT6 | 2.7424 | 9.25E-58 |
| 25 mM Fumaric acid | g2562 | ACT10084 | MEU1 | 2.7043 | 4.34E-35 |
| 25 mM Fumaric acid | g4513 | DEL15196 | MED1 | 2.6839 | 1.95E-32 |
| 25 mM Fumaric acid | g1522 | ACT03398 | MSY1 | 2.6518 | 4.27E-42 |
| 25 mM Fumaric acid | g2162 | DEL05098 | INP2 | 2.6218 | 3.23E-17 |
| 25 mM Fumaric acid | g1896 | ACT05792 | SIL1 | 2.6002 | 3.65E-84 |
| 25 mM Fumaric acid | g291 | ACT12374 | PRI1 | 2.564 | 1.33E-62 |
| 25 mM Fumaric acid | g1715 | ACT04628 | KLMA_20163 | 2.4962 | 5.00E-41 |
| 25 mM Fumaric acid | g624 | ACT25500 | JAC1 | 2.428 | 1.85E-62 |
| 25 mM Fumaric acid | g3434 | ACT15698 | MKS1 | 2.3908 | 2.56E-30 |
| 25 mM Fumaric acid | g916 | ACT27343 | TRM10 | 2.3586 | 4.65E-19 |
| 25 mM Fumaric acid | g1794 | ACT05122 | g1794 | 2.334 | 4.78E-33 |
| 25 mM Fumaric acid | g4379 | ACT21672 | KLMA_80156 | 2.2677 | 5.22E-26 |
| 25 mM Fumaric acid | g4788 | ACT24165 | GUD1 | 2.2552 | 4.16E-11 |
| 25 mM Fumaric acid | g1639 | ACT04129 | DAP2 | 2.246 | 1.76E-43 |
| 25 mM Fumaric acid | g3476 | ACT15972 | FSF1 | 2.1964 | 1.26E-33 |
| 25 mM Fumaric acid | g2753 | ACT11340 | SUR2 | 2.1823 | 3.96E-16 |
| 25 mM Fumaric acid | g680 | ACT25875 | DUG2 | 2.1479 | 1.63E-36 |
| 25 mM Fumaric acid | g3274 | ACT14660 | GCR2 | 2.0911 | 2.09E-15 |
| 25 mM Fumaric acid | g3585 | ACT16696 | VPS36 | 2.0559 | 8.25E-13 |
| 25 mM Fumaric acid | g138 | DEL01669 | TEL1 | 2.0487 | 0.0032932 |
| 25 mM Fumaric acid | g4078 | DEL13339 | HCS1 | 2.0468 | 3.13E-11 |
| 25 mM Fumaric acid | g45 | ACT22460 | PER33 | 1.8969 | 5.69E-18 |
| 25 mM Fumaric acid | g4568 | DEL15433 | RAD51 | 1.8957 | 5.64E-09 |
| 25 mM Fumaric acid | g1422 | DEL01860 | ATP22 | 1.8892 | 0.0075286 |
| 25 mM Fumaric acid | g583 | ACT25227 | CAX4 | 1.8493 | 6.35E-09 |
| 25 mM Fumaric acid | g3931 | ACT18878 | KLMA_60306 | 1.8193 | 2.19E-17 |
| 25 mM Fumaric acid | g3299 | ACT14822 | gcp | 1.7935 | 2.74E-12 |
| 25 mM Fumaric acid | g423 | ACT20770 | IRE1 | 1.7135 | 2.69E-10 |
| 25 mM Fumaric acid | g3233 | ACT14397 | SAC6 | 1.7107 | 9.46E-04 |
| 25 mM Fumaric acid | g3110 | ACT13601 | STF2 | 1.6616 | 7.01E-07 |
| 25 mM Fumaric acid | g808 | ACT26630 | MGR1 | 1.6323 | 4.64E-09 |
| 25 mM Fumaric acid | g3149 | ACT13847 | fnx1 | 1.6301 | 1.70E-08 |
| 25 mM Fumaric acid | g1859 | ACT05549 | KLMA_20313 | 1.6207 | 1.29E-07 |
| 25 mM Fumaric acid | g4091 | ACT19893 | MNN2 | 1.617 | 6.59E-06 |
| 25 mM Fumaric acid | g263 | ACT10523 | MEP3 | 1.6084 | 4.94E-10 |
| 25 mM Fumaric acid | g461 | ACT23182 | CDC9 | 1.5807 | 0.0019247 |
| 25 mM Fumaric acid | g863 | ACT26990 | UFD1 | 1.5608 | 1.02E-06 |
| 25 mM Fumaric acid | g1308 | ACT02009 | RPC31 | 1.5414 | 7.66E-10 |
| 25 mM Fumaric acid | g1493 | DEL02173 | RAG2 | 1.5385 | 0.014018 |
| 25 mM Fumaric acid | g2957 | ACT12684 | NIT3 | 1.5231 | 3.16E-08 |
| 30 mM Fumaric acid | g4399 | ACT21802 | QDR2 | 10.831 | 0 |
| 30 mM Fumaric acid | g4399 | ACT21806 | QDR2 | 10.269 | 0 |
| 30 mM Fumaric acid | g3178 | ACT14031 | UBP14 | 6.0777 | 0 |
| 30 mM Fumaric acid | g3577 | ACT16642 | FAS1 | 5.3817 | 0 |
| 30 mM Fumaric acid | g592 | ACT25286 | IMD4 | 5.2305 | 0 |
| 30 mM Fumaric acid | g4399 | ACT21801 | QDR2 | 4.785 | 0 |
| 30 mM Fumaric acid | g2142 | ACT07423 | YPR1 | 4.0725 | 6.81E-203 |
| 30 mM Fumaric acid | g3020 | ACT13106 | CSE1 | 4.0243 | 0 |
| 30 mM Fumaric acid | g1501 | ACT03264 | LYS21 | 3.8844 | 0 |
| 30 mM Fumaric acid | g1640 | ACT04144 | RPN1 | 3.5368 | 6.15E-216 |
| 30 mM Fumaric acid | g1123 | ACT00802 | NNK1 | 3.4299 | 1.93E-76 |
| 30 mM Fumaric acid | g2114 | ACT07232 | KLMA_20588 | 3.3281 | 3.09E-289 |
| 30 mM Fumaric acid | g971 | ACT27711 | KLMA_10227 | 3.2804 | 1.79E-64 |
| 30 mM Fumaric acid | g4278 | ACT21032 | GPI12 | 3.2791 | 1.98E-156 |
| 30 mM Fumaric acid | g3308 | ACT14880 | PDR5 | 3.1379 | 2.77E-155 |
| 30 mM Fumaric acid | g694 | ACT25965 | LAS21 | 3.1375 | 2.57E-93 |
| 30 mM Fumaric acid | g878 | ACT27085 | GIS2 | 3.1304 | 3.95E-53 |
| 30 mM Fumaric acid | g2445 | ACT09339 | RRN3 | 3.078 | 6.37E-69 |
| 30 mM Fumaric acid | g902 | ACT27254 | ACO2a | 2.9787 | 1.12E-39 |
| 30 mM Fumaric acid | g1538 | ACT03500 | PEX32 | 2.8542 | 1.09E-68 |
| 30 mM Fumaric acid | g1291 | ACT01896 | HGT1 | 2.854 | 3.95E-53 |
| 30 mM Fumaric acid | g292 | ACT12438 | KLMA_70317 | 2.8203 | 1.43E-40 |
| 30 mM Fumaric acid | g547 | ACT24985 | QDR3 | 2.8108 | 4.20E-226 |
| 30 mM Fumaric acid | g2227 | ACT07985 | VMA2 | 2.7874 | 4.00E-59 |
| 30 mM Fumaric acid | g4077 | DEL13335 | ATP7 | 2.7562 | 2.37E-14 |
| 30 mM Fumaric acid | g4406 | ACT21855 | ZUO1 | 2.6927 | 3.92E-61 |
| 30 mM Fumaric acid | g267 | ACT10790 | KLMA_70292 | 2.5663 | 2.43E-16 |
| 30 mM Fumaric acid | g3618 | ACT16917 | TOM1 | 2.5642 | 3.28E-33 |
| 30 mM Fumaric acid | g1345 | ACT02252 | KLMA_10605 | 2.4957 | 7.67E-49 |
| 30 mM Fumaric acid | g2525 | ACT09854 | ATP16 | 2.4669 | 4.43E-56 |
| 30 mM Fumaric acid | g987 | ACT27810 | SDS23 | 2.4595 | 8.75E-30 |
| 30 mM Fumaric acid | g3014 | ACT13062 | DUG1 | 2.4144 | 1.30E-37 |
| 30 mM Fumaric acid | g2043 | ACT06768 | PKP1 | 2.3343 | 6.77E-16 |
| 30 mM Fumaric acid | g2056 | ACT06851 | CAP2 | 2.2976 | 1.03E-16 |
| 30 mM Fumaric acid | g2189 | ACT07724 | ARG81 | 2.2743 | 3.25E-33 |
| 30 mM Fumaric acid | g1670 | ACT04334 | COX16 | 2.271 | 8.78E-12 |
| 30 mM Fumaric acid | g4307 | ACT21221 | NSE1 | 2.2326 | 3.23E-25 |
| 30 mM Fumaric acid | g659 | ACT25730 | GIP4 | 2.226 | 3.66E-30 |
| 30 mM Fumaric acid | g3813 | ACT18126 | g3813 | 2.1355 | 3.99E-24 |
| 30 mM Fumaric acid | g2521 | ACT09831 | RTG1 | 1.9937 | 2.21E-36 |
| 30 mM Fumaric acid | g1896 | ACT05792 | SIL1 | 1.9475 | 1.54E-25 |
| 30 mM Fumaric acid | g2989 | ACT12888 | SKI8 | 1.9466 | 1.28E-04 |
| 30 mM Fumaric acid | g624 | ACT25500 | JAC1 | 1.9314 | 1.22E-24 |
| 30 mM Fumaric acid | g680 | ACT25875 | DUG2 | 1.8156 | 9.72E-19 |
| 30 mM Fumaric acid | g3585 | ACT16696 | VPS36 | 1.8092 | 2.73E-07 |
| 30 mM Fumaric acid | g3700 | ACT17407 | VPS54 | 1.7763 | 2.21E-08 |
| 30 mM Fumaric acid | g407 | ACT19751 | PAC2 | 1.7106 | 4.10E-09 |
| 30 mM Fumaric acid | g3201 | ACT14193 | KLMA_40132 | 1.6967 | 1.80E-14 |
| 30 mM Fumaric acid | g291 | ACT12374 | PRI1 | 1.6857 | 2.05E-11 |
| 30 mM Fumaric acid | g3331 | ACT15031 | OCH1 | 1.6438 | 7.22E-07 |
| 30 mM Fumaric acid | g1390 | DEL01718 | SNQ2 | 1.5703 | 3.58E-04 |
| 30 mM Fumaric acid | g1522 | ACT03398 | MSY1 | 1.5204 | 3.40E-04 |
| 42 mM Fumaric acid | g4399 | ACT21802 | QDR2 | 10.972 | 0 |
| 42 mM Fumaric acid | g4399 | ACT21806 | QDR2 | 9.7333 | 0 |
| 42 mM Fumaric acid | g3178 | ACT14031 | UBP14 | 5.5647 | 0 |
| 42 mM Fumaric acid | g592 | ACT25286 | IMD4 | 4.9176 | 0 |
| 42 mM Fumaric acid | g3020 | ACT13106 | CSE1 | 4.5154 | 0 |
| 42 mM Fumaric acid | g2142 | ACT07423 | YPR1 | 3.7074 | 6.72E-107 |
| 42 mM Fumaric acid | g1501 | ACT03264 | LYS21 | 3.5609 | 2.30E-178 |
| 42 mM Fumaric acid | g3577 | ACT16642 | FAS1 | 3.5103 | 2.50E-187 |
| 42 mM Fumaric acid | g2114 | ACT07232 | KLMA_20588 | 3.0591 | 4.76E-166 |
| 42 mM Fumaric acid | g4399 | ACT21801 | QDR2 | 3.0521 | 2.82E-164 |
| 42 mM Fumaric acid | g694 | ACT25965 | LAS21 | 3.0118 | 1.39E-67 |
| 42 mM Fumaric acid | g2445 | ACT09339 | RRN3 | 2.9279 | 4.33E-48 |
| 42 mM Fumaric acid | g3308 | ACT14880 | PDR5 | 2.8817 | 2.36E-90 |
| 42 mM Fumaric acid | g1640 | ACT04144 | RPN1 | 2.8142 | 1.29E-60 |
| 42 mM Fumaric acid | g902 | ACT27254 | ACO2a | 2.8104 | 7.51E-27 |
| 42 mM Fumaric acid | g4278 | ACT21032 | GPI12 | 2.6555 | 4.85E-50 |
| 42 mM Fumaric acid | g1123 | ACT00802 | NNK1 | 2.6533 | 4.36E-19 |
| 42 mM Fumaric acid | g878 | ACT27085 | GIS2 | 2.6403 | 8.58E-21 |
| 42 mM Fumaric acid | g2056 | ACT06851 | CAP2 | 2.1818 | 1.50E-11 |
| 42 mM Fumaric acid | g987 | ACT27810 | SDS23 | 2.0943 | 6.90E-13 |
| 42 mM Fumaric acid | g267 | ACT10790 | KLMA_70292 | 2.0179 | 1.35E-04 |
| 42 mM Fumaric acid | g971 | ACT27711 | KLMA_10227 | 2 | 2.33E-05 |
| 42 mM Fumaric acid | g3014 | ACT13062 | DUG1 | 1.938 | 5.92E-13 |
| 42 mM Fumaric acid | g1538 | ACT03500 | PEX32 | 1.8557 | 2.94E-09 |
| 42 mM Fumaric acid | g4406 | ACT21855 | ZUO1 | 1.8501 | 7.77E-11 |
| 42 mM Fumaric acid | g4077 | DEL13335 | ATP7 | 1.8171 | 0.0078687 |
| 42 mM Fumaric acid | g659 | ACT25730 | GIP4 | 1.7964 | 7.60E-11 |
| 42 mM Fumaric acid | g1291 | ACT01896 | HGT1 | 1.7945 | 7.03E-06 |
| 42 mM Fumaric acid | g3618 | ACT16917 | TOM1 | 1.7391 | 2.93E-05 |
| 42 mM Fumaric acid | g3813 | ACT18126 | g3813 | 1.7037 | 3.10E-08 |
| 42 mM Fumaric acid | g2525 | ACT09854 | ATP16 | 1.7002 | 9.27E-11 |
| 42 mM Fumaric acid | g624 | ACT25500 | JAC1 | 1.6952 | 1.06E-12 |
| 42 mM Fumaric acid | g292 | ACT12438 | KLMA_70317 | 1.6842 | 0.0012576 |
| 42 mM Fumaric acid | g2521 | ACT09831 | RTG1 | 1.6183 | 3.91E-14 |
| 42 mM Fumaric acid | g1670 | ACT04334 | COX16 | 1.5654 | 0.006548 |
| 42 mM Fumaric acid | g1345 | ACT02252 | KLMA_10605 | 1.5508 | 6.87E-06 |
| 42 mM Fumaric acid | g2227 | ACT07985 | VMA2 | 1.5303 | 6.18E-04 |
| 42 mM Fumaric acid | g4307 | ACT21221 | NSE1 | 1.5167 | 2.51E-04 |
